# ELP-dependent expression of MCL1 promotes resistance to EGFR inhibition in triple-negative breast cancer cells

**DOI:** 10.1101/2020.03.29.014894

**Authors:** Peter Cruz-Gordillo, Megan E. Honeywell, Thomas Leete, Michael J. Lee

## Abstract

Targeted therapies for the treatment of cancer are generally thought to exploit oncogene addiction, a phenomenon in which a single oncogene controls both the growth and survival of the tumor cell. Many well-validated examples of oncogene addiction exist; however, the utility of oncogene targeted therapies varies substantially by cancer context, even among cancers in which the targeted oncogene is similarly dysregulated. For instance, epidermal growth factor receptor (EGFR) signaling can be effectively targeted in EGFR-mutant non-small cell lung cancer (NSCLC), but not in triple-negative breast cancer (TNBC), where EGFR is activated to a similar degree. We find that EGFR controls a similar signaling/transcriptional network in TNBC and EGFR-mutant NSCLC cells, but only NSCLC cells respond to EGFR inhibition by activating cell death. To address this paradox and identify mechanisms that contribute to insensitivity to EGFR inhibition in TNBC, we performed a genome-wide CRISPR-Cas9 genetic knockout screen. Our screen identifies the Elongator (ELP) complex as a mediator of insensitivity to EGFR inhibition in TNBC. Depleting ELP proteins caused high levels of apoptotic cell death, in an EGFR inhibition-dependent manner. We find that the tRNA-modifying function of the ELP complex promotes drug insensitivity, by facilitating expression of the anti-apoptotic protein MCL1. Furthermore, pharmacological inhibition of MCL1 synergizes with EGFR inhibition across a panel of genetically diverse TNBC cells. Taken together, we find that TNBC “addiction” to EGFR signaling is masked by the ELP complex, and our study provides an actionable therapeutic strategy to overcome this resistance mechanism by co-targeting EGFR and MCL1.

**One sentence summary:** The Elongator Protein (ELP) Complex masks TNBC oncogene “addiction” to EGFR signaling, by promoting expression of the anti-apoptotic protein MCL1.

## INTRODUCTION

Many modern therapies target signaling proteins that are mutated, amplified, or otherwise dysregulated in cancer cells. These targeted therapies are generally thought to exploit a phenomenon called “oncogene addiction”. This terminology was coined based on the observation that for many cancers, the regulation of growth and survival appears to be aberrantly coordinated through the activity of a single oncogene *(1, 2)*. Oncogene addiction is generally thought to result from the disordered regulatory circuitry in cancer cells *(2)*. Many notable examples exist in which selectively inhibiting an oncogenic kinase results in striking clinical responses. However, these responses are rarely durable and extremely variable *(3)*. For instance, while ~50% of BRAF mutant melanomas respond to BRAF inhibition, response rates are less than 5% in BRAF mutant colorectal cancers *(4)*. Factors that account for this variability between cancer types are generally not well understood, and the mechanisms underlying oncogene addiction remain unclear.

Triple-negative breast cancer (TNBC) is an aggressive subtype of breast cancer, defined the lack of expression of the three receptors that are best characterized to drive breast cancer tumorigenesis: estrogen receptor (ER), progesterone receptor (PR), and the human epidermal growth factor receptor 2 (HER2). Due to the lack of these targetable oncogenes, TNBCs are generally treated only with genotoxic chemotherapies, and most patients fail to achieve a complete response *(5)*. Some evidence does suggest that EGFR signaling may be aberrantly activated in TNBC. TNBCs generally have high levels of EGFR expression and activity *(6*, *7)*. Although EGFR is rarely mutated in TNBC, aberrant activity is caused by loss of downstream phosphatases or receptor amplification *(8*–*10)*. In spite of the observed relationship between EGFR activity and TNBC growth, drugs that inhibit EGFR signaling are consistently inefficacious in this setting *(7)*. In contrast, non-small cell lung cancers (NSCLC) harboring activating mutations in EGFR often respond to EGFR inhibitors *(11*, *12)*. Clear differences exist in the mechanisms by which EGFR is activated in TNBC versus NSCLC; however, it remains unclear what accounts for the varied responses to EGFR inhibition in different disease settings.

In this study we explored mechanisms by which TNBC cells evade EGFR targeted therapies. We find that the degree of EGFR activity and EGFR-dependent gene expression is similar between TNBCs and *bona fide* EGFR-dependent NSCLC cells, such as PC9. In contrast to PC9, which die when exposed to EGFR inhibitors, TNBC cells fail to activate cell death following exposure to erlotinib, an EGFR-specific inhibitor. To identify genes involved in the non-responsiveness of TNBC cells to EGFR targeted therapies, we performed a genome-wide genetic knockout screen using CRISPR-Cas9 mediated genome editing. Our screen revealed that the ELP complex promotes insensitivity to EGFR inhibitors in TNBC. We find that the ELP complex insulates TNBCs from erlotinib-mediated cell death, by promoting expression of the anti-apoptotic protein MCL1. Knocking down ELP complex genes promotes hyper-sensitivity to EGFR inhibitors through loss of MCL1 protein expression and activation of apoptotic cell death. Additionally, directly inhibiting MCL1 using the small molecule inhibitor S63845 synergistically enhances responses to EGFR inhibitors in TNBC cells. Taken together, this study establishes a new mechanism by which TNBC cells evade cell death following EGFR inhibition, and provides an actionable strategy for improving responses by directly targeting MCL1.

## RESULTS

### TNBC cells are insensitive to EGFR inhibition despite high-levels of EGFR signaling

NSCLC cells that harbor activating mutations in EGFR generally respond to EGFR inhibitors *(11*, *12)*. In contrast, TNBCs are generally insensitive to EGFR inhibition, despite having high levels of EGFR expression *(7*, *13)*. To more closely investigate subtype-specific differences in responses to EGFR inhibitors, we profiled sensitivities to the EGFR inhibitor, erlotinib, in a panel of TNBC cell lines. Erlotinib was tested across an eight-point, log10 drug titration, and drug sensitivity was measured using a SYTOX green-based assay *(14)*. Because these cell lines grow at different rates, we scored drug sensitivity using the normalized Growth Rate inhibition value (GR). GR values facilitate a more accurate comparison of drug responses across cell lines by removing artefactual differences in drug response that are due to differences in the growth rate between cells *(15)*. We compared erlotinib responses in TNBC cells to drug sensitivity observed in PC9, an EGFR-mutant NSCLC cell line that is known to be sensitive to EGFR inhibition *(16)*. As expected, we found that erlotinib strongly inhibited growth in PC9 cells, with GR values near or below zero at high doses (Fig. 1A). Responses were more varied in TNBC cells. Complete resistance was observed at all doses for three of six TNBC cells profiled, whereas intermediate responses were observed for the other three TNBCs (Fig. 1A). In the TNBC cells with intermediate responses to erlotinib, the maximum responses observed were in the positive portion of the GR scale (0.3 - 0.7), indicating net population growth even at the highest concentrations of erlotinib tested.

**Fig. 1:**
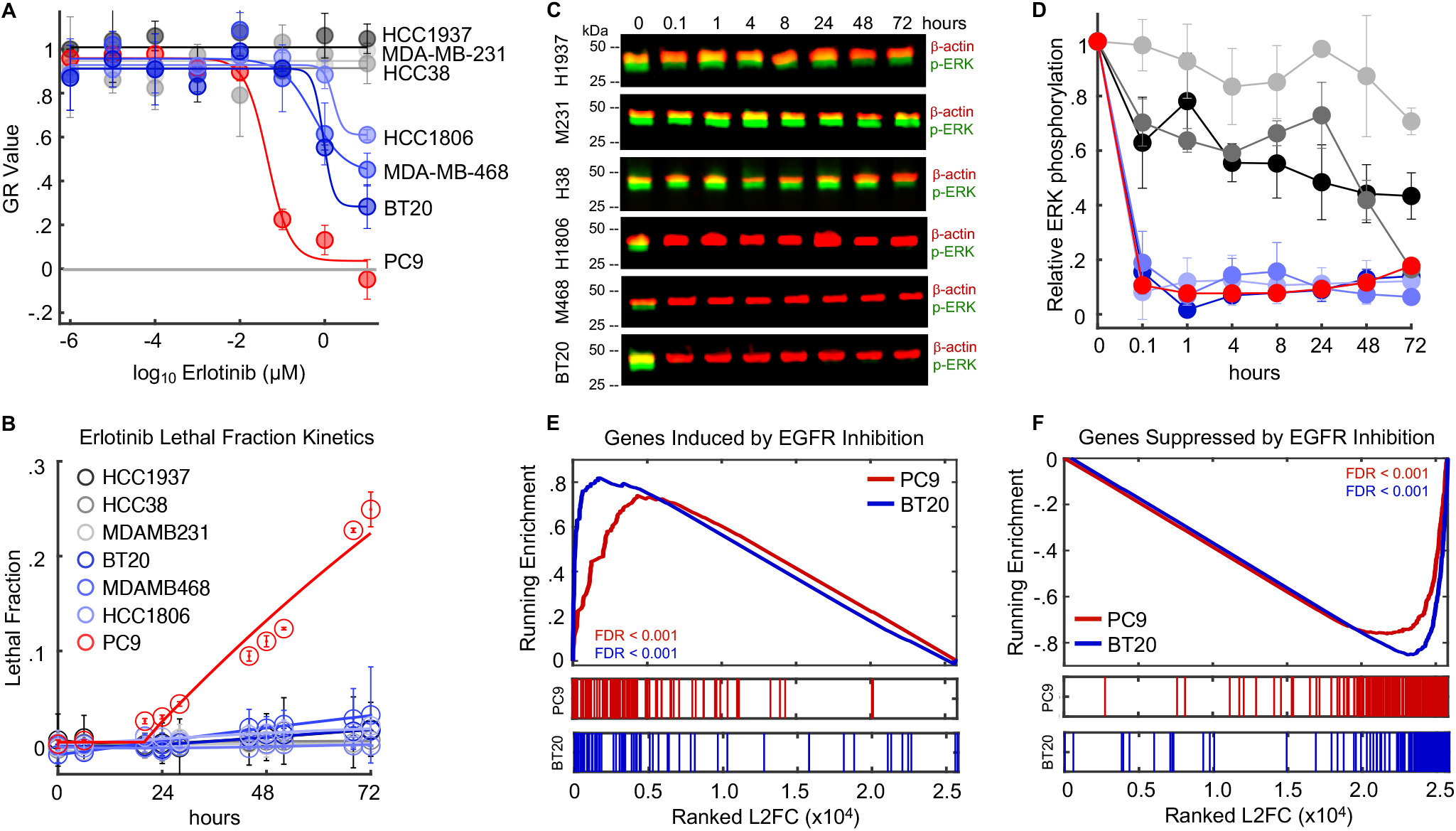
TNBC cells are insensitive to EGFR inhibition despite high-levels of EGFR dependent signaling. **(A-B)** Erlotinib sensitivity in TNBC cells evaluated using a SYTOX green based death assay. **(A)** Normalized Growth Rate Inhibition value (GR) computed after 72-hour exposure to erlotinib. PC9 (red), is a NSCLC cell line with an activating mutation in EGFR. Other cells are TNBCs that did not respond to erlotinib (grey) or had partial responses (blue). Data are the mean +/− SD from 3 biological replicates. **(B)** Erlotinib-induced Lethal Fraction Kinetics. Data are mean +/− SD from 3 biological replicates. **(C-D)** ERK phosphorylation (p-ERK) following 10 μM erlotinib exposure, monitored as a downstream EGFR dependent signal. Data in (C) are representative of 3 independent biological replicates. See also Fig. S1-2. **(D)** p-ERK quantified from western blots in (C). Data are mean +/− SD from 3 biological replicates. Data in (D) are colored as in panel (A). **(E-F)** Gene Set Enrichment Analysis (GSEA) of mRNA expression in PC9 or BT20 cells given erlotinib for 24 hours. Rank ordered log2 Fold Change (L2FC) for treated vs. untreated samples used to evaluate genes that are induced (E) or depleted (F) by EGFR inhibition in EGFR-driven NSCLC cells. Gene signatures used in comparison are from msigDB (‘Kobayashi EGFR Signaling 24-hr UP’ in (E) and ‘Kobayashi EGFR Signaling 24-hr DN’ in (F)).

The GR value reports drug response in terms of the net population growth rate, but for any given value, it is unclear to what extent the observed response is due to growth arrest, cell death, or both. Thus, we also measured drug-induced lethal fraction (*e.g.* dead cells divided by total cells), which more specifically reports drug-induced cell killing *(14*, *17)*. Evaluation of lethal fraction kinetics revealed significant drug-induced cell death in PC9, but not in any of the TNBCs tested (Fig. 1B). We also computed drug GRADE, a metric that precisely scores the degree to which cell death contributes to an observed drug response *(18)*. The drug GRADE for erlotinib in PC9 cells was 45 (*e.g.* 45% of the decrease in population size is caused by cell death, Fig. S1) and erlotinib-induced cell death was observed at low doses. In contrast, GRADEs for erlotinib in TNBC cells were much lower, ranging from 0 - 23% (Fig. S1). Furthermore, erlotinib-induced death was not observed in any TNBC cells, except at high doses. Thus, erlotinib exposure results in high levels of cell death in EGFR-driven PC9 cells, but only partial growth inhibition in TNBCs.

Given that erlotinib-induced growth inhibition in TNBC was only partial, we suspected that higher concentrations of the drug may be required to fully inhibit EGFR signaling. To inspect this, we measured phosphorylation of ERK, a critical downstream kinase that drives growth factor induced proliferation *(19)*. For TNBC cells that were completely unresponsive to erlotinib, we observed that ERK activity was also unchanged, even at the highest doses of erlotinib that we tested (Fig. 1C). Unexpectedly, however, for TNBCs that responded to erlotinib, we observed that ERK signaling was completely inhibited in spite of the observed partial response (Fig. 1C-D and Fig. S2). Furthermore, the degree, kinetics, and duration of ERK inhibition were similar in these TNBC cells to what we observed in PC9 (Fig. 1D). Thus, in many TNBC cells, erlotinib appears to be an effective EGFR inhibitor, but these cells continue to proliferate without EGFR/ERK signaling.

One possible explanation for these observations is that EGFR controls a different signaling/transcriptional network in EGFR-driven NSCLC cells when compared to TNBC or other cell types. Indeed, in the context of oncogene addiction, it is often suggested that the coordinated control of growth and death by a single protein is the result of aberrant network circuitry *(2*, *20)*. To test this, we performed RNA-seq in BT20 cells, a TNBC cell line with intermediate sensitivity to erlotinib. We compared these data to a previously published dataset of gene expression changes in PC9 following EGFR inhibition *(21)*. In both cases, data were collected for untreated cells and following 24-hour exposure to erlotinib. We analyzed erlotinib-induced changes in gene expression using gene set enrichment analysis (GSEA) *(22)*. We focused on a previously annotated signature of gene expression changes that were observed in EGFR-driven NSCLC cells when exposed to an EGFR inhibitor *(23)*. For these genes that were previously shown to be induced or depleted in EGFR-mutant NSCLC cells treated with EGFR inhibitors, we observed similar changes in PC9 and BT20 cells (Fig. 1E-F). Thus, although these cells are derived from different types of cancer, with different EGFR aberrations, the signaling network controlled by EGFR in these two cell lines appears to be similar.

### Genome-wide screen using CRISPR-Cas9 mediated genome editing reveals that the ELP complex contributes to erlotinib insensitivity in TNBC

TNBC cells are consistently insensitive to EGFR inhibition, in spite of the similarities between TNBC and EGFR-mutant NSCLC cells in EGFR-dependent signaling and gene regulation. To identify genes that may be contributing to erlotinib insensitivity in TNBC we performed a genome-wide single gene knockout screen. We used the GeCKO v2 pooled sgRNA library, which has six sgRNAs targeting each gene in the genome, and 1000 non-targeting sgRNA controls *(24)*. We infected spCas9-expressing BT20 cells at low MOI, with a coverage of ~300x per sgRNA. Cells were either left untreated or treated with 10 μM erlotinib for roughly 3-4 population doublings. To identify genes that contribute to erlotinib insensitivity we focused on genes that drop out of the library in erlotinib treated cells compared to untreated cells. A consensus “best” method for analyzing this type of screen has not yet emerged. The best performing method in any given scenario likely depends on experiment-specific parameters, which affect the distributions of targeting/non-targeting sgRNAs and the magnitude of changes within the data being analyzed *(25)*. Thus, rather than selecting an analysis method *a priori,* we tested several analysis strategies in parallel *(26*–*30)*. Collectively, these strategies tested different methods for determining: sgRNA-level fold change, gene-level scores from the distribution of sgRNA fold changes, and statistical significance within the gene-level data. To evaluate the quality of each analysis stream, we determined the ability of each method to score core essential genes in the comparison of untreated cells to the “T0” input library (Fig. S3) *(31)*.

Based on this evaluation, we selected a conservative and straight-forward approach that was similar to prior methods, such as drugZ *(32)* (Fig. 2A and Fig. S3). Briefly, we included all recovered sgRNAs (*e.g.* no trimming), and computed fold-changes at the sgRNA level using DESeq2 *(26)*. We determined gene-level scores by computing the mean of the sgRNAs targeting each gene. These scores were then z-scored relative to the mean and standard deviation of the non-targeting controls. To determine statistical cut-offs, an empiric p-value was determined from z-scored fold-changes by bootstrapping based on the sgRNA data. Empiric p-values were then FDR corrected for statistical robustness.

**Fig. 2:**
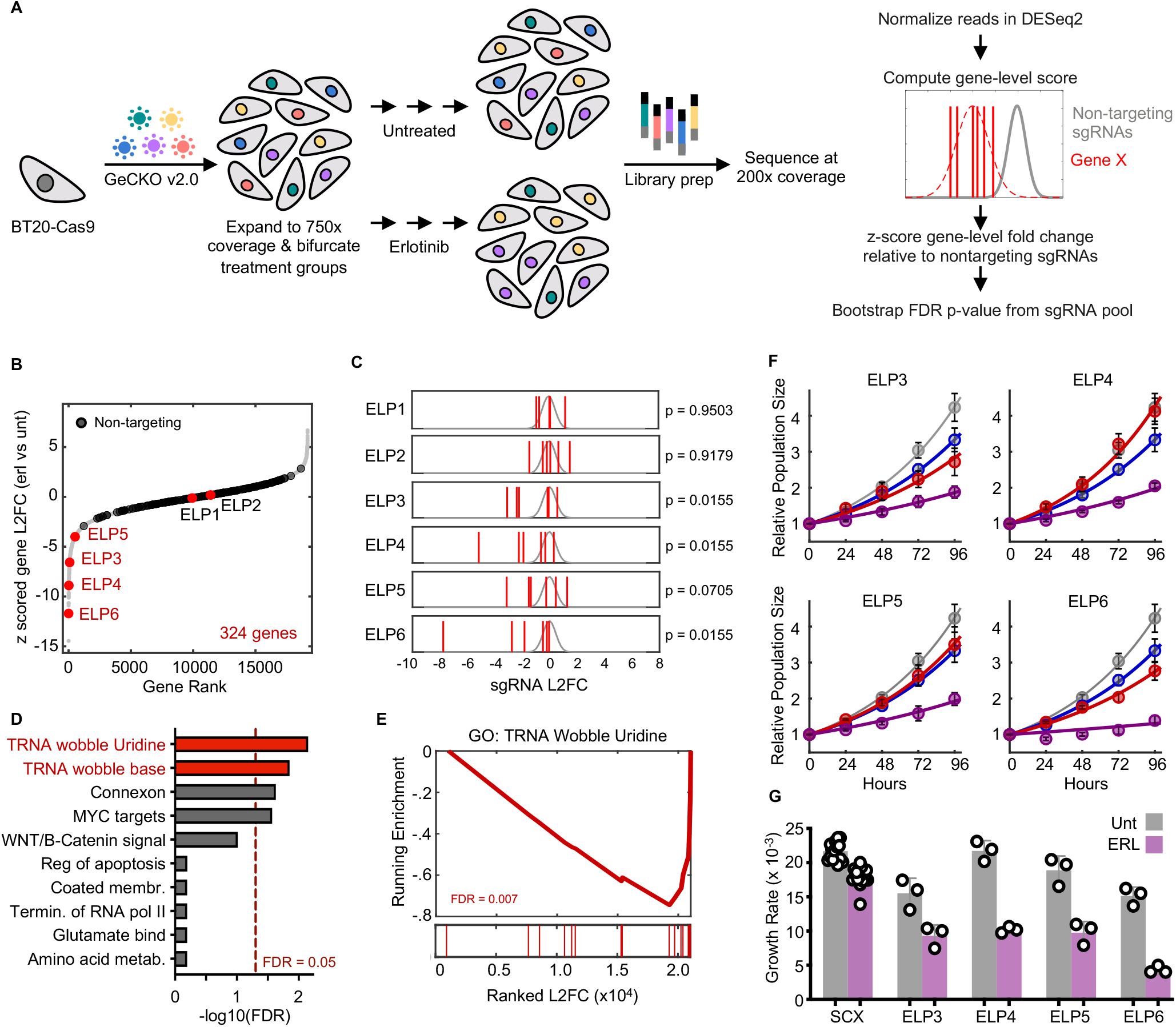
Genome-wide screen using CRISPR-Cas9 mediated genome editing reveals that the ELP complex contributes to erlotinib insensitivity in TNBC. **(A)** Schematic overview of CRISPR screen. See also Fig. S3. **(B-C)** Fold change in recovery of BT20 cells harboring gene knockouts when comparing erlotinib treated to untreated cells. **(B)** 324 genes were differentially recovered. Six ELP complex genes are highlighted (red), and non-targeting “genes” highlighted in black. Data are the z-scored log_2_ fold change (L2FC) of gene-level data. **(C)** sgRNA-level data for the six ELP genes. FDR corrected p-values shown. Grey curve is the distribution of all sgRNAs; red lines are the six individual sgRNAs for a given ELP gene (non-z scored). **(D-E)** GSEA of the CRISPR screening data. **(D)** Ten most enriched signatures within msigDB. FDR cut-off for significance shown. **(E)** Example signature shown for the GO term: TRNA Wobble Uridine. FDR p-value shown. **(F-G)** ELP validation using quantitative microscopy. **(F)** Automated image analysis used to score population growth rate for BT20 cells with or without an ELP targeting siRNA. Grey: scrambled RNA untreated. Blue: scrambled RNA + 10 μM erlotinib. Red: ELP targeting siRNA pool. Purple: ELP targeting siRNA pool + 10 μM erlotinib. See Fig. S4 for knockdown validation. **(G)** Growth rates of curves in (f). Data are population growth per hour. The growth rate of untreated BT20 cells was ~ 43 hours (0.023 increase per hour, e.g. 23 × 10^−3^). Data in (F) and (G) are mean +/− SD for 16 biological replicates in SCX and 3 biological replicates in ELP siRNA conditions.

Our analysis strategy identified 324 genes that were differentially recovered in cells exposed to erlotinib when compared to untreated cells, with 295 of these genes being depleted in erlotinib treated cells (Fig. 2B). Among the most depleted genes were four of the six components of the Elongator complex (ELP1-6, Fig. 2B-C). The ELP complex was first identified based on physical association with RNA polymerase II *(33*, *34)*. Although ELP proteins do play roles in histone acetylation and transcription, studies reveal that the central function for the ELP proteins is in modifying a subset of tRNAs in the U34 wobble base position. ELP-dependent modifications at this site are required for function of these tRNAs *(35)*. Additionally, U34 modifying enzymes have recently been found to contribute to drug resistance in BRAF mutant melanomas and ER+ breast cancers *(36*, *37)*.

To distinguish between the transcriptional and U34-modifying functions of the ELP complex, we analyzed our gene knockout screen data using GSEA. Only 4 gene signatures within the molecular signatures database were significantly enriched or depleted within our data, with the top two signatures being the GO annotations “tRNA wobble uridine modification” and a similar signature “tRNA wobble base modification” (Fig. 2D-E). Enrichment for these signatures was driven not only by the ELP genes, but also by depletion of the CTU1 and CTU2 enzymes. CTU1 and 2 are involved in the U34-modifying functions of the ELP complex, but not the transcriptional functions of ELP *(38)*. Thus, these data further clarify that the U34 modifying functions of the ELP complex contribute to erlotinib insensitivity in BT20 cells.

To verify the results of our screen, we knocked down expression of ELP3-6 using siRNAs. The population growth rate for each knockdown was quantified in the presence and absence of erlotinib using automated microscopy. Consistent with expectations, we found that erlotinib exposure caused a slowing of the population growth rate by roughly 20% (46 hours versus 55 hours for the doubling time, Fig. 2F-G). For all four ELP proteins tested, knocking down expression strongly enhanced population growth suppression in the erlotinib treated cells (Fig. 2F-G and Fig. S4). Notably, while knockdown of ELP3, ELP5, and ELP6 resulted in some growth slowing even in the absence of erlotinib, knockdown of ELP4 strongly suppressed growth in an erlotinib-dependent manner. These data further highlight the synergistic interaction between ELP knockdown and erlotinib exposure.

### ELP complex promotes survival of TNBC cells exposed to erlotinib by promoting expression of MCL1

Knocking down the ELP proteins enhanced responses to erlotinib; however, it was unclear if this was due to further inhibiting the growth rate of cells, increasing the death rate, or a combination of these phenotypes. In *bona fide* cases of EGFR “oncogene addiction”, EGFR inhibition causes robust cell death (Fig. 1C). Thus, we measured the drug-induced lethal fraction over time following erlotinib exposure, using a SYTOX-based cell death assay *(14)*. As expected, erlotinib did not result in cell death in triple-negative BT20 cells, even when applied at high concentrations. For all four ELPs tested, however, ELP knockdown significantly increased erlotinib-induced cell killing (Fig. 3A and Fig. S4).

**Fig. 3:**
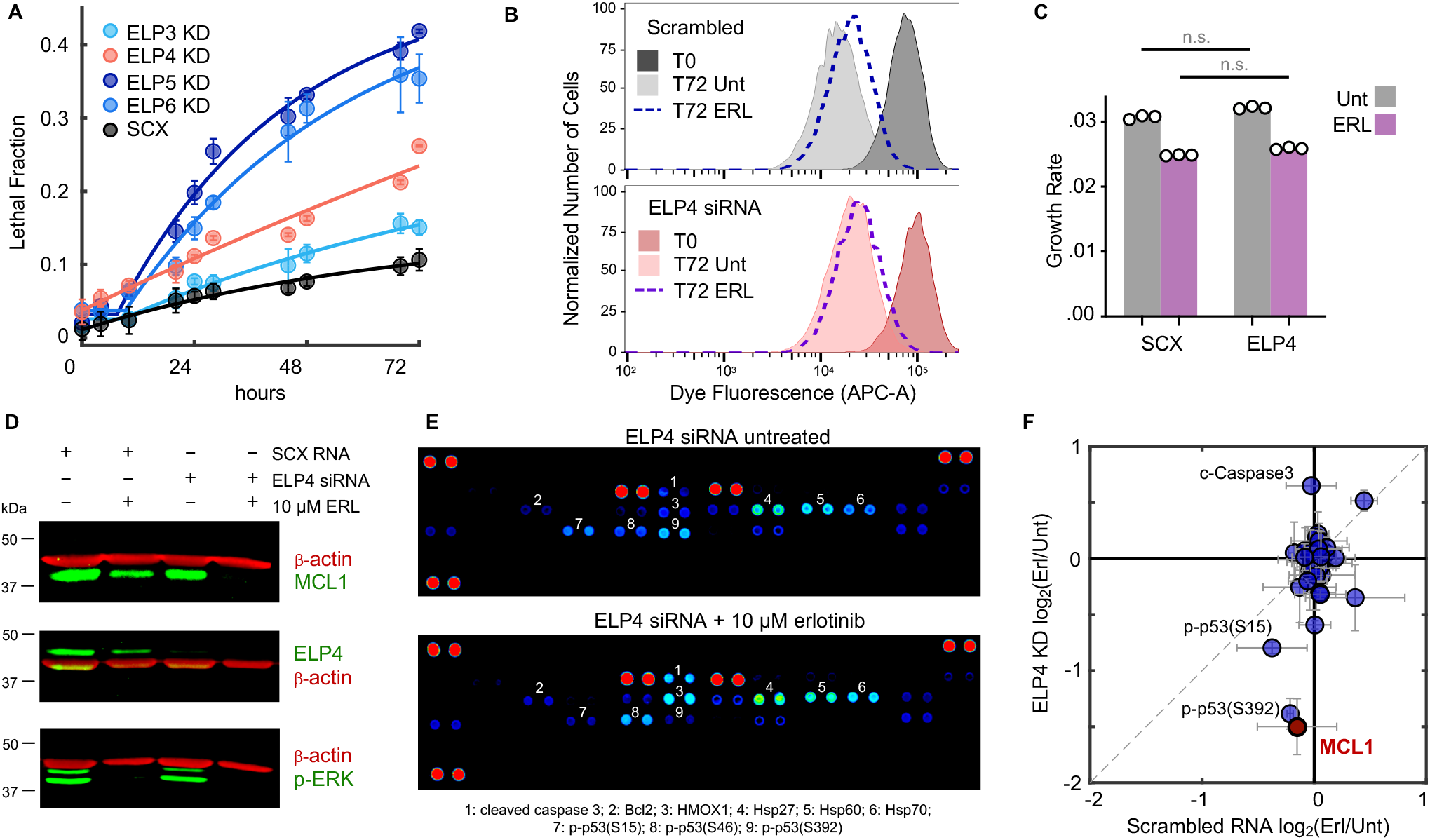
ELP complex promotes survival of TNBC cells exposed to erlotinib by promoting expression of MCL1. **(A)** Lethal fraction kinetics following application of 10 μM erlotinib. Cells tested were BT20 + scrambled RNA (SCX), or BT20 + siRNA targeting ELP3, 4, 5, or 6 (ELP KD). Data are mean +/− s.d. from three biological replicates. **(B)** Cell proliferation dye dilution to determine growth rate of live cells. BT20 cells given a scrambled RNA (top) or ELP4 targeted siRNA (bottom) were labelled with CellTrace proliferation dye. Cells were either left untreated (Unt) or given 10 μM erlotinib (ERL). Samples collected prior to drug addition or following 72 hours. Representative dye fluorescence distributions from flow cytometry shown. Data are representative of 3 independent biological replicates. **(C)** Quantification of cell growth rate from data in (b). Data are mean +/− s.d. of 3 independent biological replicates. **(D)** MCL1 protein expression in BT20 cells + SCX or + ELP4 siRNA, with or without 10 μM ERL. Data are representative of 3 independent biological replicates which all showed similar results. See also Fig. S6. **(E)** Proteome profiler apoptotic array featuring 39 apoptotic proteins. Data shown are representative of 2 independent biological replicate blots from ELP4 siRNA untreated or ELP4 siRNA + 10 μM ERL. Both blots showed similar data. See also Fig. S7. **(F)** Quantification of (d) and (e). Data are mean +/− s.d. of three (d) or two (e) replicates.

Drug-induced cell death is generally not mutually exclusive with drug-induced growth inhibition *(18)*. To determine the degree to which growth inhibition is also contributing to the observed drug response, we also measured the proliferation rate of erlotinib-treated cells following ELP knockdown. We focused on ELP4 because, unlike the other ELP members, ELP4 knockdown did not cause any population growth defects in the absence of erlotinib (Fig. 2F-G). To measure the growth rate of live cells, rather than the effective population growth rate, we calculated the rate of dilution of a cell tracking dye over time. As expected, these data reveal that the true growth rate of cells is somewhat faster than the effective population growth rate, due to baseline levels of cell death (*i.e.* true doubling time of 35 hours, compared to a population growth rate of 46 hours, Fig. 2G and 3B-C). These data also revealed that ELP4 knockdown caused a selective increase in the death rate, without altering the cell proliferation rate (Fig. 3C). Our prior drug screening found that this is a somewhat rare phenotype that is generally only observed for direct activators of cell death, such as BH3 mimetics *(18)*.

Having found that ELP4 knockdown facilitates erlotinib-induced cell death in TNBC cells, we next aimed to determine the mechanism by which the ELP complex was controlling death in these cells. The ELP complex facilitates modification of a subset of tRNAs at the U34 base. These modifications are required for decoding AAA^Lys^, GAA^Glu^, and CAA^Gln^ codons during mRNA translation *(39)*. In yeast, depletion of ELP proteins causes protein misfolding and accumulation of protein aggregates *(40)*. Additionally, in the context of cancer, inhibiting ELP function causes ribosome stalling and inefficient translation of transcripts that are enriched for U34 codons *(37)*. To identify ELP target genes that may be mediating the erlotinib insensitivity in TNBC, we sought to identify negative regulators of apoptosis whose expression was dependent on ELP complex function. We noticed in our CRISPR screen that MCL1 was one of the most depleted genes, suggesting that erlotinib insensitivity depends on MCL1 expression (12^th^ most depleted gene; z-scored L2FC = −9.7, Fig. 2 and Table S1-2). Furthermore, no other apoptotic regulatory genes were found on our list of significantly depleted genes.

MCL1 is a member of the BCL2 family, and a potent negative regulator of apoptosis *(41)*. To determine if MCL1 expression is regulated by ELP complex activity, we measured erlotinib-induced changes in MCL1 protein levels, with and without ELP4 depletion. Using quantitative immunoblotting, we found that MCL1 protein expression is strongly depleted in ELP4 knockdown cells treated with erlotinib (Fig. 3D and Fig. S5). In cells in which ELP4 was not depleted, erlotinib did not significantly alter MCL1 levels. To determine if this was unique to MCL1, or if other apoptotic proteins were also regulated by the ELP complex, we profiled a panel of well-validated apoptotic regulatory proteins using the Proteome Prolifer Apoptotic Array. These data show that very few changes are observed following erlotinib exposure in wild-type BT20 cells, consistent with our observations that erlotinib does not induce death in these cells in the presence of robust ELP activity (Fig. S6). In the context of ELP4 knockdown, however, several apoptotic proteins change in their expression (Fig. 3E-F). Notably, caspase-3 cleavage is increased in ELP4 KD cells exposed to erlotinib, confirming that the death observed is likely due to an apoptotic mechanism. Other erlotinib-induced changes include decreased phosphorylation of p53 on multiple phosphorylation sites, and increased Bcl2 expression (Fig. 3F and Fig. S6). Notably, these changes would be expected to inhibit apoptosis, and thus are not likely to promote the cell death that is observed.

### MCL1 inhibition synergistically enhances sensitivity to erlotinib in TNBC cells

Our data suggest that the ELP complex contributes to erlotinib insensitivity in TNBC cells by reinforcing expression of MCL1, a negative regulator of apoptosis. Thus, we next tested if direct inhibition of MCL1 would potentiate sensitivity to erlotinib in these TNBC cells. We tested all pairwise combinations of erlotinib and the MCL1 inhibitor, S63845 *(42)*. In BT20 cells, both drugs had very modest efficacy as single agents, killing less than 30% of all cells even at saturating high doses (Fig. 4A). When these drugs were added in combination, however, cell killing was substantially increased (Fig. 4A). To evaluate the combinatorial drug-drug interaction, we analyzed the data using the Loewe isobologram convention *(43*, *44)*. Under the Loewe dose additivity model, one would expect two drugs to produce linear isobols when added in combination (*i.e.* linear trajectory connecting similarly efficacious doses). Analysis of erlotinib-S63845 combinations in BT20 cells revealed a strongly synergistic interaction between these drugs (Fig. 4B). For instance, 1 μM S63845, an inefficacious concentration of this drug, potentiated erlotinib sensitivity by roughly 100-fold (Fig. 4C).

**Fig. 4:**
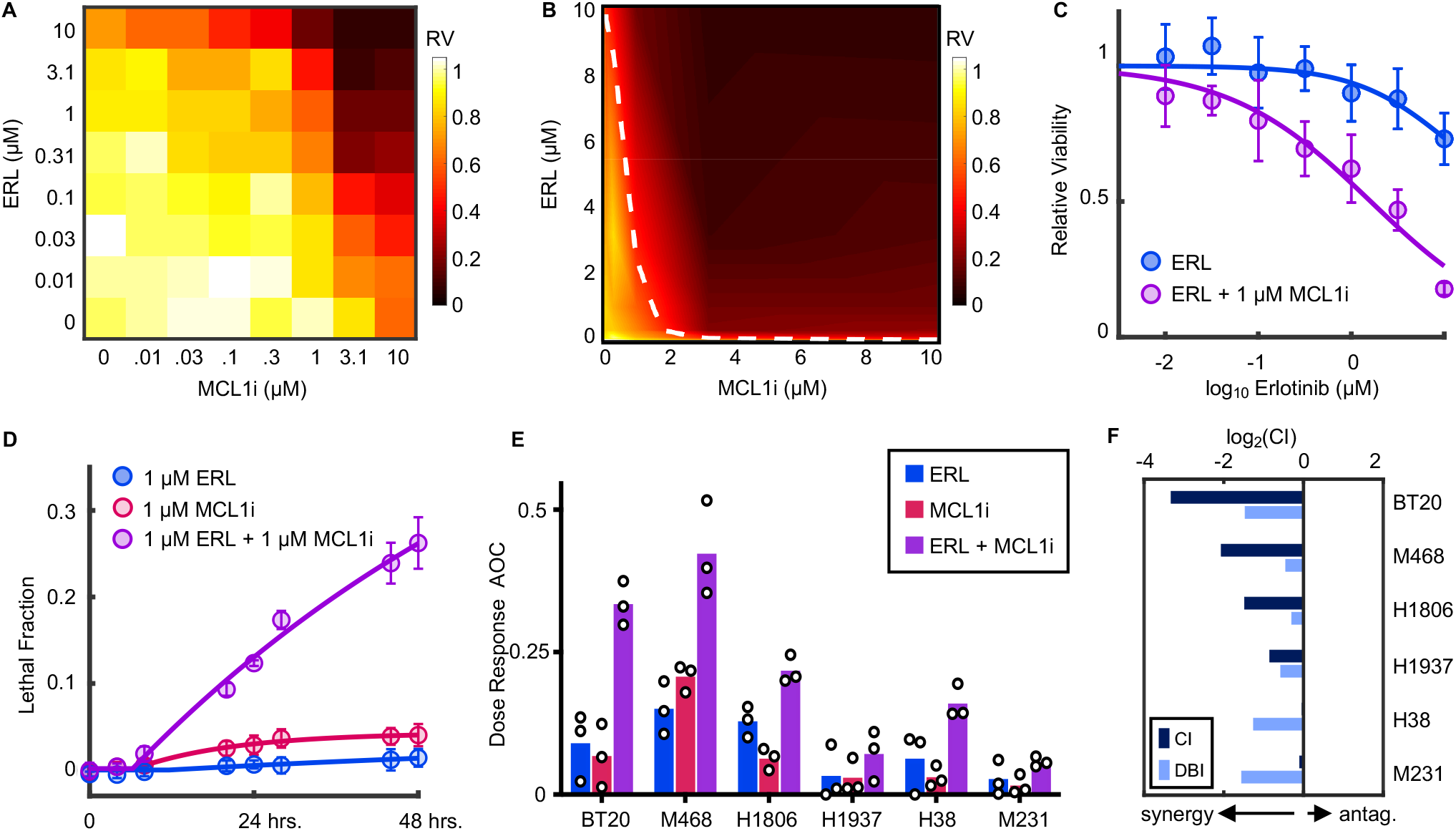
MCL1 inhibition synergistically enhances sensitivity to erlotinib in TNBC. **(A)** Full dose titration of erlotinib (ERL) and S63845 (MCLi) in BT20 cells. Data are relative viability of drug treated cells compared to untreated cells at 72 hours. Heatmap is scaled according to mean values from 3 biological replicate experiments. **(B)** Isobologram analysis for data in (A). Data are arrayed in linear scale with linear interpolation. Dashed white line represents ERL+MCLi combinations that result in 50% response (i.e. 50% isobol). **(C)** ERL sensitivity at 72 hours with or without 1 μM MCLi. As in (A), data are mean +/− s.d. from 3 independent biological replicates. **(D)** Lethal fraction kinetic responses in BT20 cells treated with 1μM ERL, 1μM MCL1i, or both. **(E)** ERL, MCL1i, or combination responses at varied doses in a panel of TNBCs. Drugs were tested as single agents or in 1:1 fixed ratio combinations across 7 doses. Data are mean +/− s.d. of area over the dose-response curve (AOC) for 3 biological replicates. **(F)** Combination Index (CI) and Deviation from Bliss Independence (DBI) computed for combinations of ERL and MCL1i in TNBC cells.

To determine if the observed drug effect was due to growth inhibition or cell killing, we also measured the drug-induced lethal fraction. As expected, erlotinib or S63845 did not result in significant cell death when added as single agents. Alternatively, combinations of erlotinib and S63845 induced high levels of cell death (Fig. 4D). Furthermore, the death onset time, death rate, and maximum lethal fraction observed were similar to what we previously observed for erlotinib sensitivity following ELP4 depletion. Additionally, these responses were also similar to the responses observed in EGFR-driven PC9 cells (Fig 1B, 3A). Thus, MCL1 inhibition, while not lethal to cells, promotes erlotinib sensitivity in a manner that is similar to that observed in the absence of robust ELP function, and similar to *bona fide* examples of EGFR oncogene addiction.

To determine if MCL1 potentiation of erlotinib sensitivity was generalizable to other TNBCs, we profiled combinations of erlotinib + S63845 in a panel of TNBC cell lines. Drugs were tested alone and in fixed dose-ratio combinations (1:1 equimolar dosing). In all TNBCs tested, sensitivity to erlotinib was improved by addition of S63845 (Fig. 4E and Fig. S7). However, the degree of improvement was variable across the six cell lines. To more formally score these drug-drug interactions, we used two well-validated conventions, the Chou-Talalay Combination Index (CI) and Deviation from Bliss Independence (DBI) *(43, 49)*. CI scores drug-drug interactions relative to a dose additivity reference model, whereas DBI scores interactions relative to a response independence reference model *(14*, *44)*. These measures of drug-drug interaction can both be used to identify synergistic or antagonistic drug combinations; however, CI and DBI tend to vary due to the differences in how additivity or independence are defined *(45)*. For interactions between MCL1 inhibition and EGFR inhibition in TNBC, scores were synergistic for all cell lines tested, by both CI and DBI measures, further highlighting the robustness of this drug-drug interaction (Fig. 4F and Fig. S7). Taken together, these data demonstrate that MCL1 promotes the insensitivity to EGFR inhibition that is commonly observed within the TNBC subtype.

## DISCUSSION

In this study we explored differences between EGFR-driven NSCLC cells and TNBC, a cancer subtype that exhibits high levels of EGFR signaling, but does not respond to EGFR inhibitors. Our genetic screen identified that the ELP complex - in particular the tRNA modifying function of the ELP complex - is important for promoting insensitivity to EGFR inhibition. We found that ELP activity insulates the cells from activating apoptotic death, by promoting expression of the anti-apoptotic protein, MCL1. Finally, we find that direct targeting of MCL1 synergistically potentiates erlotinib sensitivity in a panel of genetically-unrelated TNBC cells.

These data complement two studies that have recently highlighted a role for ELP-dependent translation in promoting cancer drug resistance. The ELP complex was found to promote resistance to BRAF inhibition in BRAF-driven melanoma, through specific stabilization of HIF1a *(37)*. Additionally, the ELP tRNA-modifying activity was found to promote resistance to ER inhibition in ER+ breast cancer, through a broad-spectrum “translational offsetting”, which stabilizes ER-dependent protein expression, despite the loss of ER-dependent mRNAs *(36)*. Our finding that ELP activity also drives the characteristic EGFR-insensitivity of TNBCs further extends the reach of ELP-dependent drug resistance. Furthermore, at least for TNBC cells, our study highlights a targetable protein, MCL1, which can reverse ELP-dependent drug resistance.

A question that remains unclear is why MCL1 protein levels are particularly sensitive to the tRNA modifying activity controlled by the ELP complex. MCL1 mRNA is a direct substrate of the U34 tRNAs that require ELP-dependent modifications. MCL1 has 18 ELP-sensitive codons and loss of U34 tRNAs may lead to difficulty translating these codons *(37)*. However, considering the length of the MCL1 mRNA, 18 ELP-sensitive codons is roughly average for the human genome (~ 51 ELP sensitive codons per 1000 codons on average). Thus, it remains unclear why MCL1 protein expression was strongly dysregulated, but not the expression of many other apoptotic regulatory proteins. It is possible that in addition to MCL1, other key MCL1 regulatory proteins are also direct ELP targets. MCL1 has a relatively short protein half-life, due to an amino-terminal PEST sequence that promotes protein degradation *(46)*. Protein turnover is further regulated by phosphorylation of MCL1 by a variety of MAPKs, which slows the protein turnover rate *(47)*. Additionally, MCL1 mRNA expression and protein stability are controlled by PI3K/mTOR signaling in a variety of different contexts *(48)*. Thus, the ELP complex may be promoting MCL1 protein expression by regulating multiple substrates, including MCL1 itself, and potentially other proteins that contribute to MCL1 turnover, growth factor signaling, MAPK signaling, and/or PI3K/mTOR signaling. Thus, a major benefit of our finding is that regardless of which ELP substrate(s) are functionally coordinated by ELP activity to regulate MCL1 levels, direct targeting of MCL1 pheno-copies ELP deletion, leading to enhanced sensitivity to EGFR inhibition. This finding provides a strong rationale for the development of combination drug therapies involving co-inhibition of EGFR and MCL1 in TNBC, and potentially other subtypes of cancer.

More broadly, our study reveals an important new insight into the phenomenology of “oncogene addiction”. TNBC cells would not typically qualify as being “addicted” to EGFR, as these cells do not respond to loss of EGFR signaling. Our data show that when compared to NSCLC cells with EGFR-activating mutations, EGFR in TNBC signals at a similar level, activates a similar set of genes, and is equally inhibited by commonly used EGFR inhibiting drugs. Thus, EGFR plays the same role in many TNBC cells as in NSCLC cells that are “addicted” to EGFR. This reality is masked by the activity of the ELP complex. Oncogene addiction has been the conceptual motivation for the development and use of targeted therapies for more than twenty years. Despite considerable effort and focused research in this area, relatively few cancer subtypes respond in the striking manner observed in the best case examples *(3)*. This study highlights a new mechanism by which phenotypes associated with oncogene addiction can be masked. In future studies it will be interesting to determine the degree to which the ELP complex contributes to drug resistance in other settings, and if MCL1 inhibition is broadly useful for potentiating targeted therapies outside of the TNBC subtype.

## MATERIALS AND METHODS

### Cell lines and general reagents

BT20, MDA-MB-468, HCC1806, HCC1937, HCC38, and MDA-MB-231 cells were obtained from the American Type Culture Collection (ATCC). PC-9 (NSCLC) cells were a generous gift from J. Pritchard (Penn St.). All cells were maintained at low passage numbers (less than 20 passages from the original vial). BT20 cells were grown in Minimum Essential Medium (ThermoFisher Scientific, CAT# 11090081). MDA-MB-468 and MDA-MB-231 cells were grown in Dulbecco’s Modified Eagle Medium (DMEM, Fisher Scientific, CAT# MT10017CV). PC9, HCC1806, HCC1937, and HCC38 cells were grown in Roswell Park Memorial Institute (RPMI)-1640 medium (ThermoFisher Scientific CAT# 11875119). In all cases, base medium was supplemented with 10% FBS(ThermoFisher Scientific, CAT# SH30910.03, LOT# SH40014-13), 2 mM glutamine (ThermoFisher Scientific CAT# MT25005CI), and penicillin/streptomycin (ThermoFisher Scientific, CAT# MT30002CI).

Erlotinib hydrochloride salt was purchased from LC Laboratories (CAT# E-4007). S63845 (MCL-1 inhibitor) was purchased from Selleck Chemicals (CAT# S8383). Anti-Phospho-p44/42 MAPK (ERK1/2—Thr202/Tyr204, CAT# 9101) and anti-MCL-1 (D35A5 rabbit anti-human primary, CAT# 5453) antibodies were purchased from Cell Signaling Technology. Anti-ELP4 (rabbit anti-human primary, CAT# NBP2-16322) antibody was purchased from Novus Biologicals. Monoclonal anti-β-actin (mouse anti-human primary, CAT# A2228) antibody was purchased from Sigma-Aldrich. Proteome Profiler Human Apoptosis Array kit (CAT# ARY009) and Pepstatin A aspartic protease inhibitor (#1190/10) were purchased from R&D Systems. The cOmplete protease inhibitor cocktail (CAT# 11697498001) and PhosSTOP phosphatase inhibitor tablets (CAT# 4906845001) were purchased from Millipore Sigma. The siGENOME non-targeting siRNA control pool was purchased from Horizon Discovery (CAT# D-001206-14-05). Human ELP3 (ENTREZ# 55140, CAT# M-015940-01-005), human ELP4 (ENTREZ#26610, CAT# M-016927-01-005), human ELP5 (ENTREZ# 23587, CAT# M-017992-02-005), and human ELP6 (ENTREZ# 54859, CAT# M-020705-01-005) siRNA were each purchased as a SMARTpool of 4 siGENOME siRNAs targeting different areas of the gene from Horizon Discovery.

### Lethal fraction kinetic analysis using a fluorescence plate reader

Lethal fraction (LF) kinetics were determined as describe previously *(14)*. SYTOX Green (ThermoFisher Scientific, CAT# S7020) is a nuclear marker that can only enter cells following death and loss of membrane integrity. Fluorescence of dead cells was monitored with a Tecan Spark microplate reader using an excitation/emission of 503/524. SYTOX was used at a final concentration of 5 μM, and a gain was selected for each cell line which achieved linearity below the saturation limit of the detector. SYTOX-based assays were performed in optical bottom black-walled plates (Corning, CAT# 3904), with 2500-5000 cells seeded in each well depending on each cell line’s growth rate. Cells were plated in an initial volume of 90 μL of media. Drugs and SYTOX were diluted to 10X final concentration in phosphate buffered saline (PBS), and 10 μL of each drug/drug combination were added to each well at the start of the experiment. Fluorescence readings were taken at the indicated timepoints for each experiment, with measurement frequency optimized to capture the onset time, rate, and maximum death achieved by each drug. At the end of each experiment, cells were permeabilized with the addition of 0.15% Triton X-100 (Fisher, CAT# BP151-500) and incubation at 37°C for >1.5 hours. Detergent induced cell permeabilization at the assay endpoint allows for the determination of total cell number, a crucial number for calculating LF. Additionally, for kinetic experiments, an untreated plate of cells was lysed at the time of drug addition. LF was then determined at each timepoint using pre- and post- permeabilization numbers for each plate.

### Quantitative Immunoblotting

For generation of protein lysates for immunoblotting, cells were seeded at 1 - 1.5 million cells in 10-cm dishes and allowed to adhere overnight. Erlotinib (ERL) was added at t = 0 hours at a final concentration of 10 μM, and cellular lysates were prepared at the indicated time points. Briefly, media was removed by aspiration, and plates were washed two times with 2 mL of ice-cold PBS. Cells were lysed by adding 400 μL of Sodium dodecyl sulfate (SDS)-lysis buffer (50 mM Tris-HCl, 2% SDS, 5% glycerol, 5 mM EDTA, 1 mM NaF, 10 mM β-GP, 1 mM PMSF, 1 mM Na_3_VO_4_, and a protease inhibitor and phosphatase inhibitor tablet each). Lysates were collected with cell scrapers and centrifuge spin-filtered through 0.2 μm multi-well filters to remove DNA (Pall, CAT# 5053). After filtration, each lysate concentration was determined by the Pierce BCA Protein assay kit, according to the manufacturer’s instructions (ThermoFisher Scientific, CAT# 23225). Lysate concentrations were normalized to 0.5 mg/mL for SDS-PAGE loading. Samples were run either on pre-cast E-PAGE 8% 48-well gels (ThermoFisher Scientific, CAT# EP04808) or in hand-poured 8% SDS-PAGE gels. Gels were subsequently transferred using a semi-dry iBlot fast gel transfer system (ThermoFisher Scientific) on nitrocellulose membranes (ThermoFisher Scientific, CAT# IB301031). Membranes were then blocked in a 50% PBS and 50% Odyssey Blocking Buffer (ThermoFisher Scientific, CAT# 927-40000) solution, on a rocking shaker for 1 hour at room temperature. Membranes were then incubated overnight on a rocking shaker at 4°C in primary antibody (diluted 1:1000 in a 50% PBS-0.1%Tween and Odyssey blocking buffer solution). The following morning, membranes were incubated in primary anti-β-actin antibody (diluted 1:15000 in a 50% PBS-0.1%Tween and Odyssey blocking buffer solution) for 1 hour on a rocking shaker at room temperature. Following two, 5-minute washes with PBS-0.1%Tween, membranes were incubated in secondary antibodies (diluted 1:15000 in a 50% PBS-0.1%Tween and Odyssey blocking buffer solution) conjugated to infrared dyes (LICOR, IRDye 680RD, goat anti-mouse IgG secondary, CAT# 926-68070; IRDye 800CW, goat anti-rabbit IgG secondary, CAT# 926-32211) for 1 hour at room temperature on a rocking shaker, washed 3 times, and stored in PBS. Blots were then visualized using a LICOR Odyssey CLx scanner. Protein expression signal levels were quantified with LICOR’s ImageStudio software. Expression levels were individually normalized to β-actin loading controls and all samples normalized to a BT-20 serum shock control.

### Knockdown using SMARTpool siRNAs

Knockdown experiments were performed according to the manufacturer’s suggestions. Briefly, cells were seeded in full media at 350,000 cells per well in 6-well dishes and allowed to adhere overnight. Lipofectamine RNAiMAX (ThermoFisher Scientific, CAT# 13778075) was diluted 1:100 in Opti-MEM reduced serum medium (ThermoFisher Scientific, CAT# 31985088). The siGENOME non-targeting control siRNA pool or the ELP-complex gene siGENOME SMARTpools were stored at stock concentrations of 20 μM at −80°C. For transient transfections, each reagent was diluted 1:100 in Opti-MEM reduced serum medium. The diluted lipofectamine RNAiMAX and siGENOME pools were mixed 1:1 (250 μL of each) and incubated at room temperature for 15 minutes. The 500 μL mixture of RNAiMAX+siGENOME pools were then added drop-wise to the cells (1:5 dilution) for a final concentration of 20 nM for the siRNA pools (5 nM per each siRNA within the pool). Transiently transfected cells were incubated at 37°C (and 5% CO_2_) for 24 hours before being re-plated with fresh media for subsequent experiments. Drug treatment began 48 hours after the initiation of transfection, to ensure efficient knockdown.

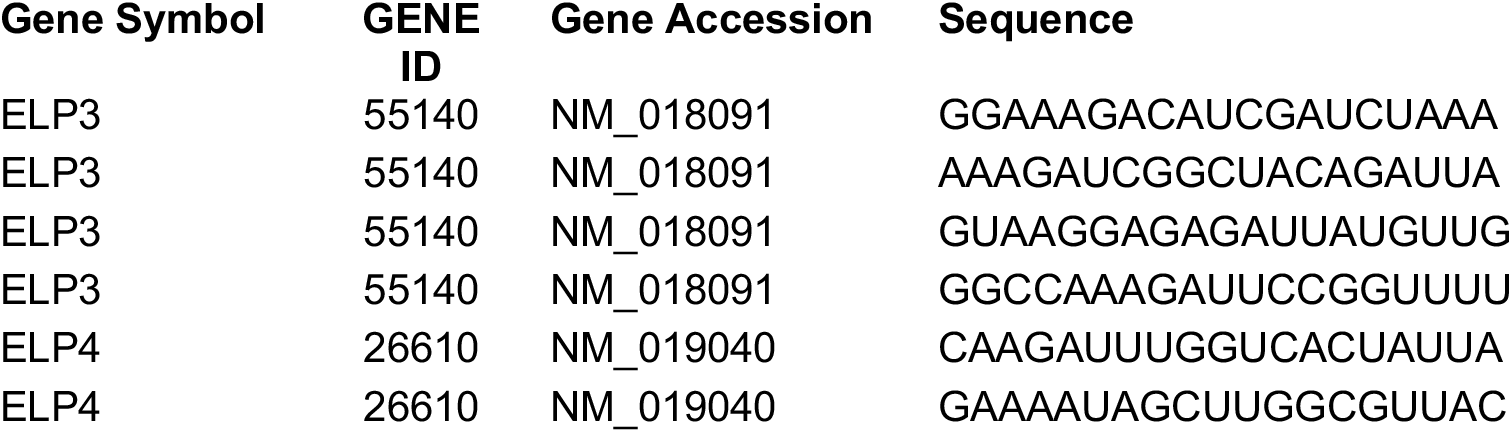
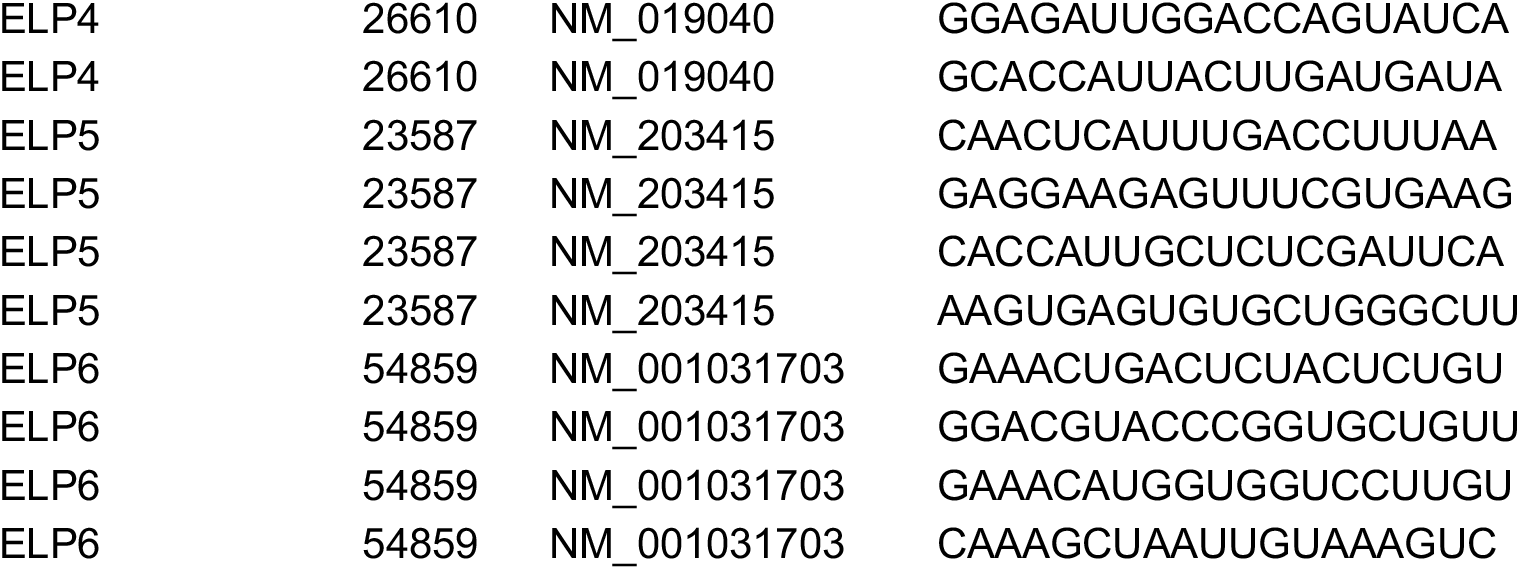

### Quantitative-PCR and qPCR primers

Following transient transfections of ELP genes and non-targeting controls, BT20 cells were seeded at 500,000 cells per 10cm plate and allowed to adhere overnight. The next day (48 hours after transfection), cells were washed twice with 2 mL of PBS. Then, 250 μL of QIAzol (CAT# 79306, Qiagen) was added to the 10 cm plate and allowed to wash the full surface of the plate. Cell lysate was collected, vortexed for 1 minute, and incubated for 5 minutes at room temperature before being snap-frozen with liquid nitrogen and stored at −80°C. Total RNA was extracted using the manufacturer’s instructions in the QIAGEN RNeasy kit (CAT# 74104, Qiagen). Each sample was at least 100 ng/μL in baseline concentration, with 260/280 ratios between 2.03-2.04 and 260/230 ratios between 1.9-2. For the reverse transcription PCR, 1 μg of total RNA (per sample) was mixed with 1 μL of Oligo-DT (ThermoFisher Scientific, CAT# 18418012), 1 μL of 10 mM dNTPs, and water for a total volume of 16.75 μL. The samples were exposed to 65°C for 5 minutes before cooled on ice quickly and spun down. Separately, a master mix of 1 μL of RNAse OUT (ThermoFisher Scientific, CAT# 10777019), 2 μL of 10x RT buffer, and 0.25 μL of reverse transcriptase (M-MuLV Reverse Transcriptase kit from New England Biolabs, CAT# M0253S) was prepared and 3.25 μL of this master mix was added to each of the samples for a total volume of 20 μL. The samples underwent the RT-PCR using the following thermocycling conditions: 90 minutes at 42°C, 5 minutes at 65°C, and then stored at −20°C. The cDNA was dilute 1:10 in water before proceeding with the quantitative PCR. Two microliters of diluted cDNA was added to a master mix of 10 μL of SYBR Green (ThermoFisher Scientific, CAT# 4385612), 1 μL each of 10 μM forward and reverse primer, and 6 μL of water, for a total reaction volume of 20 μL. The thermocycling conditions were as follows: Initial denaturation (1 cycle—10 minutes at 95°C), cycling stages (40 cycles—15 seconds at 95°C then 1 minute at 60°C with fluorescence recording at the end of each cycle), followed by a melt curve (1 cycle—1 minute at 60°C then a 0.3°C/s ramp down from 95°C with continuous fluorescence recording). The primer sets (25 nanomole DNA Oligos purchased from Integrated DNA Technologies) for the quantitative PCR were designed to skip an intron and with amplicon sizes between 70-200 nucleotides. They are as follows:

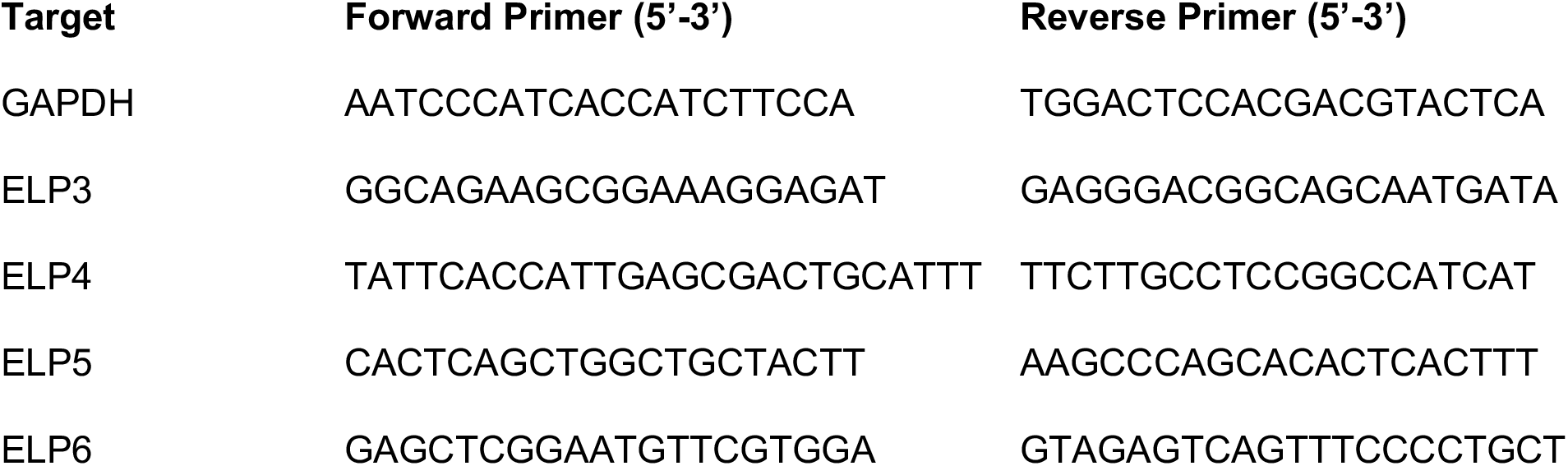

Quantitative PCR was performed in technical triplicate. After the quantitative PCR, the delta-delta CT (ΔΔCT) method was used to determine the siRNA mediated on-target knockdown of the ELP complex mRNAs, relative to the mRNA levels of these genes in a non-targeting control genetic background. Briefly, 1) GAPDH sample CTs were subtracted from ELP gene or non-targeting control CTs, and 2) the ELP gene CTs are then subtracted from the non-targeting control CTs to obtain the ΔΔCT value per ELP gene target set. We then computed the knockdown percentage for each target ELP gene by 100*(1-(1/(2^−ΔΔCT^)).

### Gene expression analysis by RNA-seq

For analysis of erlotinib-induced changes in gene expression, BT20 cells were seeded at 300,000 cells per well in a 6-well plate and allowed to adhere overnight. The next day cells were treated with either DMSO (final concentration of 0.1%) or erlotinib (final concentration of 10 μM). After 24 hours, cells were washed twice with 1 mL of PBS. 150 μL of QIAzol was added to each well and allowed to cover the full surface of the plate. The cell lysate was collected, vortexed for 1 minute, and incubated for 5 minutes at room temperature before being snap-frozen with liquid nitrogen and stored at −80°C. Total RNA was extracted using the manufacturer’s instructions in the QIAGEN RNeasy kit. Each sample was at least 50 ng/μL in baseline concentration, with 260/280 ratios of 1.9-2.03 and 260/230 ratios between 1.9-2. RNA-seq library construction was prepared using the TruSeq RNA library preparation kit v2 (CAT# RS-122-2001, Illumina) and subsequently sequenced at a depth of at least 15 million reads per sample. The experiment was completed with biological triplicate samples per condition. For comparison to PC9 cells, publicly available RNA-seq data was used (GEO accession #GSM2692587) *(21)*. Both sets of raw sequencing reads were processed using the UMass Medical School Biocore’s DolphinNext (v1.1.10) pipeline. First, the sequencing reads were de-multiplexed and assessed for sequence quality using the barcode splitter and trimmer functions of the FASTX toolkit (v0.0.14) as well as FastQC (v0.11.8). Bowtie2 was used to count and filter out RNAs that are not relevant for our analysis (i.e., rRNAs, miRNAs, tRNAs, piRNAs, etc.). RSEM (v1.3.1) was used to align RNA-seq reads to reference transcripts and estimates gene expression levels. Hisat2 (v2.1.0) and Tophat2 (v2.1.1) align the reads to the human genome (GRCh38). DESeq2 uses a parametric fit to compute a log_2_(fold-change) and FDR-adjusted *p-value* from RSEM expected counts for each comparison (erlotinib versus DMSO-treated cells).

### GSEA

Gene-Set Enrichment Analysis (GSEA)*(22)* was performed using GSEA v4.0.3 [build: 23]. For analysis of EGFR signaling signatures in BT20 and PC9 cells, we used ranked log_2_(fold-change) of mRNA expression levels in BT20 or PC9 cells, in comparisons of erlotinib versus DMSO-treatment. The GSEA pre-ranked tool was used to analyze signature enrichment or depletion. We analyzed enrichment for previously annotated gene signatures describing genes that change in expression upon EGFR inhibition in EGFR-driven NSCLC cells (Kobayashi_EGFR_signaling_24hr_UP and DOWN). Significance was determined using 1000 gene set permutations. GSEA of Genome-Wide CRISPR-Cas9 screen hits was performed as described above using rank ordered gene-level fold change values. All gene expression signatures within msigdb were tested.

### Genome-wide CRISPR screen and analysis

CRISPR screen was performed in BT20 cells using the GeCKO v2 two vector system (Addgene, CAT# 1000000049). Both A and B libraries were amplified according to the distributor, and virus was generated using 293T cells. The lentiCas9-Blast plasmid from the GeCKO system was used to generate BT20-Cas9 cells. Following blasticidin selection, Cas9-expressing cells were transduced with the sgRNA library at a MOI of < 0.3. Cells were transduced to achieve a final library coverage of 750x after puromycin selection. A T0 sample was harvested prior to drug addition, and cells were then bifurcated into DMSO (vehicle) and Erlotinib (10 μM) treatment groups. Treatment was carried out for 6 days, sub-culturing as necessary while maintaining a library coverage >200x (approximately 3-4 population doublings). Two replicates were collected for each treatment group and frozen. gDNA was isolated from the cell pellets using a phenol-chloroform based extraction method, and sgRNA sequences were amplified by PCR. A second round of PCR was used to add multiplexing barcodes, and each gel-purified library was sequenced.

For analysis of sequenced libraries, reads were first de-multiplexed using the barcode splitter and trimmer functions of the FASTX toolkit. edgeR was used to map reads from each condition to the GeCKO library. Counts were split between guides with identical sequences, and highly duplicated sgRNA sequences (>6) were removed. The sequencing depth of each sample was normalized to the distribution of nontargeting guides. A log_2_(fold-change) was determined for each comparison of interest using a parametric fit in DESeq2. The 1000 nontargeting sgRNA controls were then randomly assigned to set of 6 “genes”. Guide-level scores were then consolidated to a single gene-level fold-change by taking the mean. Gene fold-change values were then z-scored based on the distribution of non-targeting guides. Empiric p-values were calculated by bootstrapping guide-level scores, and final FDR corrected scores were determined using the Benjamini-Hochberg procedure. The method described above was selected from a larger set of methods tested in parallel, featuring different methods of trimming guides prior to gene level fold-change calculation, and different methods to estimate gene-level scores from sgRNA level fold change values. In total, 15 unique analysis pipelines were tested in parallel (see also Supplementary Figure 3). The results from these pipelines were also compared to the outputs from other published analysis methods including MAGeCK, BAGEL, and drugZ*(27*–*29)*. The default parameters were used for analysis in drugZ. In BAGEL, the lists of essential and non-essential genes generated by Hart et al. were used as training sets*(31)*. MAGeCK was performed using nontargeting guides as a negative control.

### Growth rate measurement using live cell microscopy

Following transient transfections with ELP and non-targeting controls siRNA (described above), BT-20 cells were seeded at 5000 cells per well in a 96-well optical-bottom plate and allowed to adhere overnight. Forty-eight hours after transfection cells were treated in technical triplicate with DMSO (final concentration of 0.1%) or erlotinib (final concentration of 10 μM). While being incubated at 37°C in an IncuCyte automated microscope, wells were imaged every 24 hours for 96 consecutive hours. Four quadrants were imaged per well at 10x magnification with phase-contrast live-cell microscopy. Phase-contrast masks for identifying cell bodies were analyzed using the IncuCyte analysis software to obtain well-confluence percentages for each well at each timepoint. Well-confluence percentages were normalized to timepoint 0 to obtain relative population sizes at each timepoint. These population sizes were then fit to obtain population growth curves (2^b*x^, with b signifying the growth rate and x signifying the timepoint).

### Growth rate measurement using cell dyes

To determine the growth rate of live cells following drug exposure, BT-20 cells were washed once with 5 mL of PBS and, subsequently, stained at 37°C for 20 minutes with 1 μM (final concentration) of the CellTrace Far Red reagent (CAT# C34572, ThermoFisher Scientific). After incubation, 5 mL of complete culture medium was added, the solution was mixed and incubated at 37°C for 5 minutes. Cells were then seeded at 150,000 cells per well 6-well dishes and allowed to adhere overnight. 48 hours after transient transfection, the drug-treated plates were treated with either DMSO (final concentration of 0.1%) or erlotinib (final concentration of 10 μM) and incubated at 37°C. At each timepoint samples were harvested after trypsinization, stained with 500 nM SYTOX green for 30 minutes, and placed in a solution of 1% BSA in PBS for flow cytometry analysis. At least 10,000 live cell events were collected (SYTOX green negative) per condition. After 72 hours, treated cells were harvested, stained, and analyzed by flow cytometry. The mean fluorescence intensity (MFI) of the cell population was quantified using FlowJo. The MFIs of each sample were then fit to obtain population growth curves (2^b*x^, with b signifying the growth rate and x signifying the timepoint) and growth rates.

### Proteome Profiler based analysis of apoptotic proteins

Proteome Profiler Apoptosis Array was performed as suggested in the manufacturer’s instructions. 200,000 BT-20 cells were used per condition. Cells were collected and lysed according to instructions. Lysates were diluted to 400 ug/mL and 250 μL of the lysate was loaded per array. Blot arrays were visualized using a LICOR Odyssey CLx scanner and analyte expression was quantified with the LICOR ImageStudio software.

### Data analysis and statistics

All data analysis was performed in MATLAB unless otherwise noted. Curve fitting for drug dose response and LF kinetics were performed as described previously, using custom MATLAB code*(14)*. GSEA was performed using the GSEA 4.0.3 package, and associated graphs were generated in MATLAB. Immunoblot analysis was performed using ImageStudio v4.0.21. Drug-drug interaction scores were calculated using custom code as described previously*(14)*. Drug GRADE calculations were performed as described previously*(18)*. Flow cytometry analysis was performed using FlowJo v10.5.3. PCA was performed following z-scoring, in MATLAB using the built-in function ‘pca’.

## ACKNOWLEDGEMENTS

We thank current and past members of the Lee lab, T. Fazzio, S. Hainer, and all members of PSB for their helpful comments and critiques during the design and execution of this study. We also thank M. Walhout for her helpful comments during the preparation of this manuscript. PC9 cells were a kind gift from Justin Pritchard (Penn State Univ.).

## AUTHOR CONTRIBUTIONS

This project was conceived by PCG and MJL. All experiments were performed by PCG, with help from TL during the design and execution of the CRISPR screen. Flow cytometry analysis, quantitative immunoblotting, GSEA, and growth rate analyses were performed by PCG and MJL. Curve fitting, isobologram analysis, and drug-drug interaction were performed by MEH. Analysis of the genome-wide CRISPR screen was performed by MEH and TL. Drug GRADE and PCA were performed by MJL. Manuscript was written and edited by PCG, MEH, and MJL.

## DATA AND MATERIALS AVAILABILITY

All raw data from this study are either supplied in the Supplementary materials or will be made available upon request. RNA-seq data for BT20 with and without 10 μM erlotinib are deposited onto the Gene Expression Omnibus (accession numbers supplied when they are available). A raw counts table for the genome wide screen of BT20 cells untreated or treated with erlotinib is available in Supplementary Table 1. Fully analyzed data from our screen is available in Supplementary Table 2.

## SUPPLEMENTARY MATERIALS

**Supplementary Table 1: Raw counts from genome-wide screen of erlotinib treated BT20 cells.** Raw counts from sequencing for the genome-wide CRISPR screen described in Fig. 2. GeCKO v2 library ID (lib), gene symbol (gene), and counts data for cells treated with erlotinib (ERL_rep1 and ERL_rep2), or vehicle control (DMSO_rep1 and DMSO_rep2). The counts prior to drug addition are also included (T0).

**Supplementary Table 2: Gene-level fold-change data from genome-wide screen of erlotinib treated BT20 cells.** Fully analyzed data from the genome-wide CRISPR screen described in Fig. 2. Data provided for gene depletion rank (Rank), gene symbol (Gene), gene description (Description), FDR adjusted p-value (FDR), log2-Fold Change (L2FC), and mean counts (MeanCounts). See also Supplementary Fig. 3 and the Methods for detailed descriptions of the analysis strategy.

**Fig. S1:**
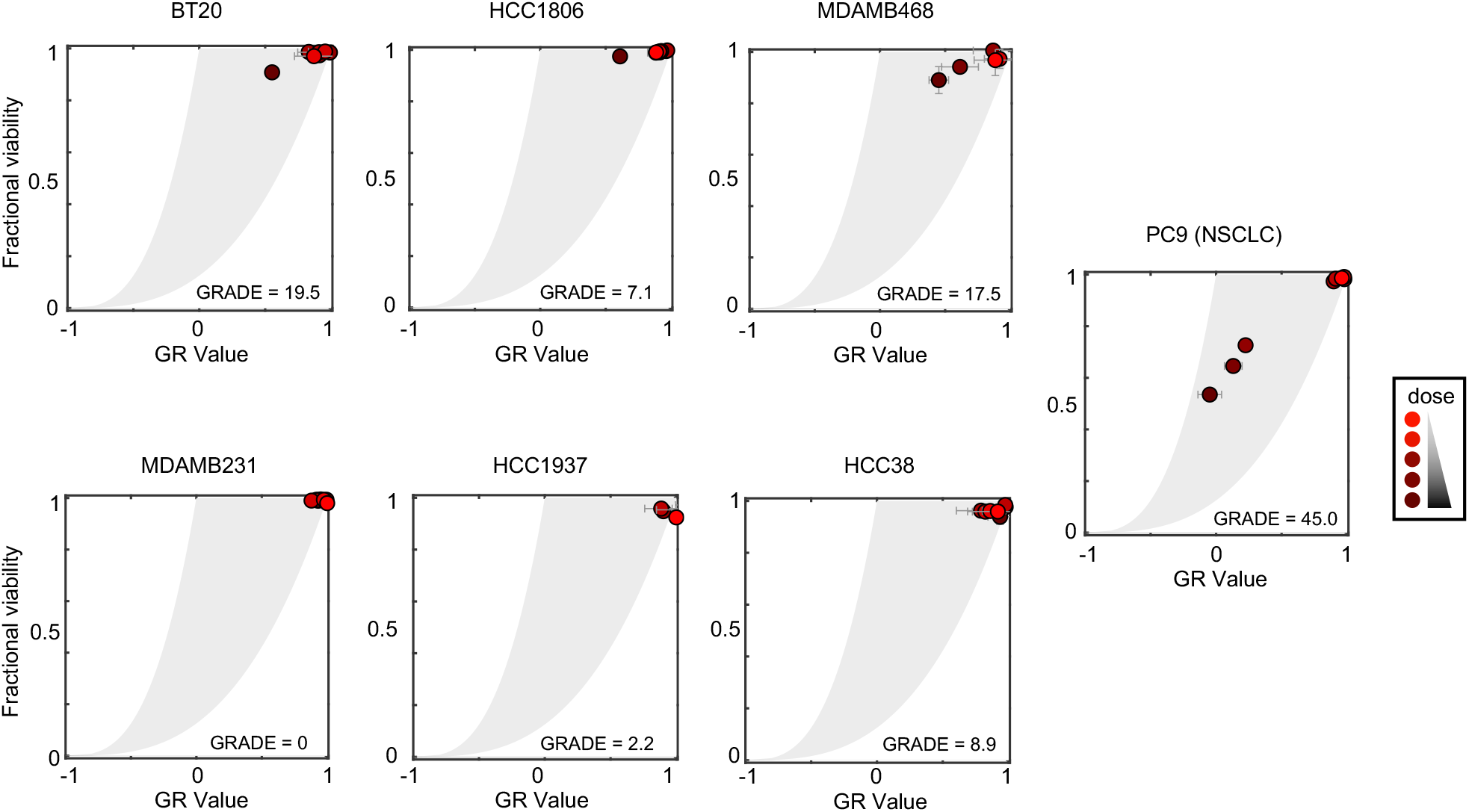
Erlotinib Drug GRADE in TNBC compared to PC9 cells. For each cell line in this study, drug GRADE (Growth Rate Adjusted DEath fraction) computed across 8 doses. Each dot represents a different dose, with darker shades used for higher doses of erlotinib (8 half-log dilutions from 0 - 10 μM). GRADE reports the fraction of the observed response that is due to cell death. Data are mean +/− SD of three biological replicates.

**Fig. S2:**
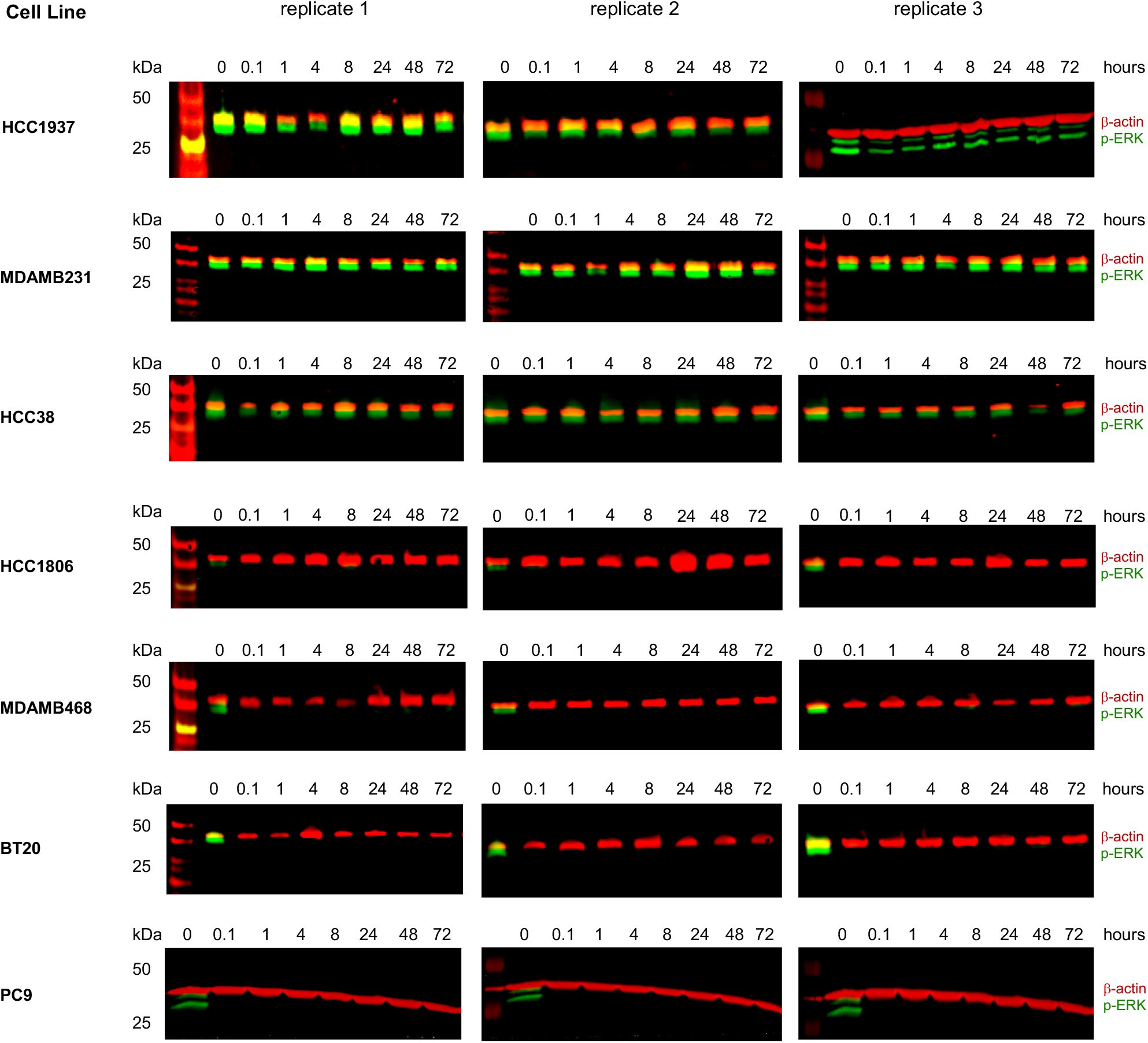
ERK activation following erlotinib treatment in TNBC cells. ERK phosphorylation (p-ERK) following 10 μM erlotinib exposure, monitored as a downstream EGFR dependent signal. Three biological replicates per cell line shown. See also Fig. 1C.

**Fig. S3:**
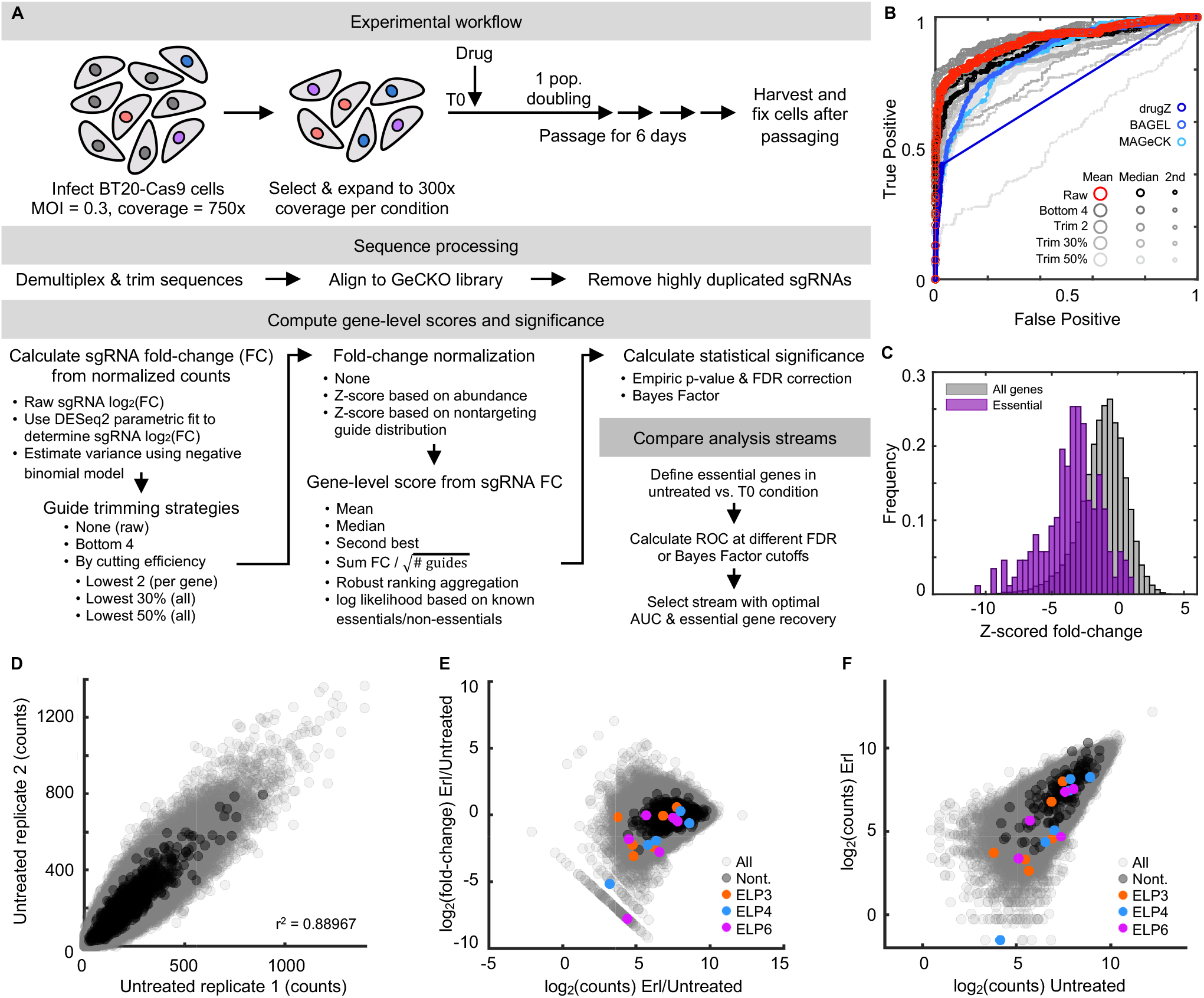
Genome-wide screen to identify genes that contribute to erlotinib insensitivity in TNBC cells. **(A)** Screen and analysis workflow. Critical steps highlighted, including methods to compute sgRNA level fold change, which sgRNAs to include, gene-level scores from sgRNA scores, statistical significance, and strategy to determine optimal analysis methods. **(B)** ROC curves for evaluation of various analysis strategies described in (A). Untreated cells compared to the T0 input control used, and data report recovery of core essential genes (True Positives) compared to non-targeting genes (False Positives) at varied FDR cut-offs. Analysis strategy selected for this study was to include all sgRNAs (no trimming, i.e. Raw), and collapse to gene level by computing the mean of sgRNAs for a given gene (red curve). ROC curve used here as the number of essential genes is approximately equivalent to the number of non-targeting “genes”. **(C)** Histogram of z-scored gene level fold changes for all genes (grey) or core essential genes from Hart et al. (purple). **(D-F)** sgRNA-level data for comparisons of erlotinib treated versus untreated cells. **(D)** sgRNA level counts for biological replicates. Pearson’s correlation coefficient shown. **(E)** sgRNA-level counts versus fold change observed in treated/untreated. **(F)** Scatter plot of sgRNA level counts for treated and untreated samples. See also Fig. 2 for gene-level data.

**Fig. S4:**
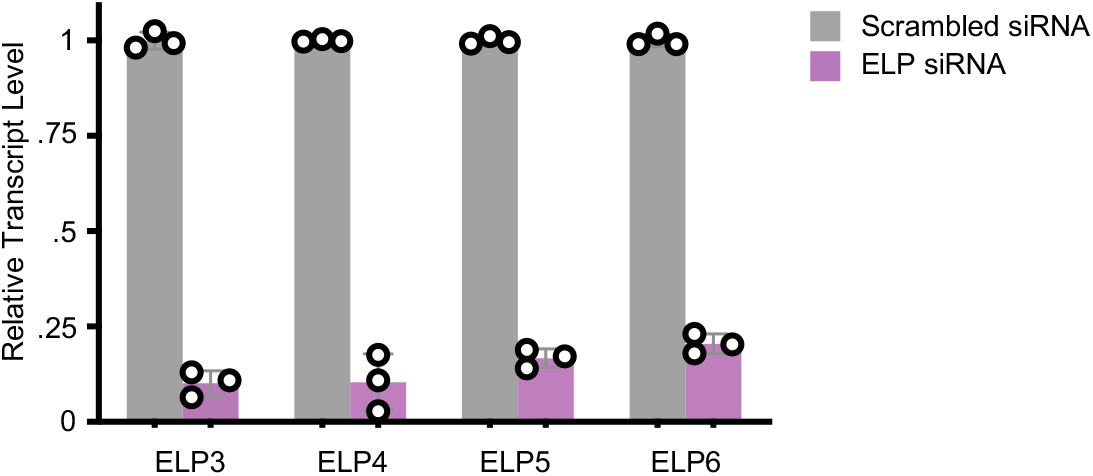
Validation of ELP knockdown by pooled siRNAs. qPCR performed to determine transcript levels for ELP3-6 following knockdown by siRNA. Data are mean +/− s.d. of three replicates. See also Fig. 3 and Fig. S5 for validation of ELP4 using Western blot.

**Fig. S5:**
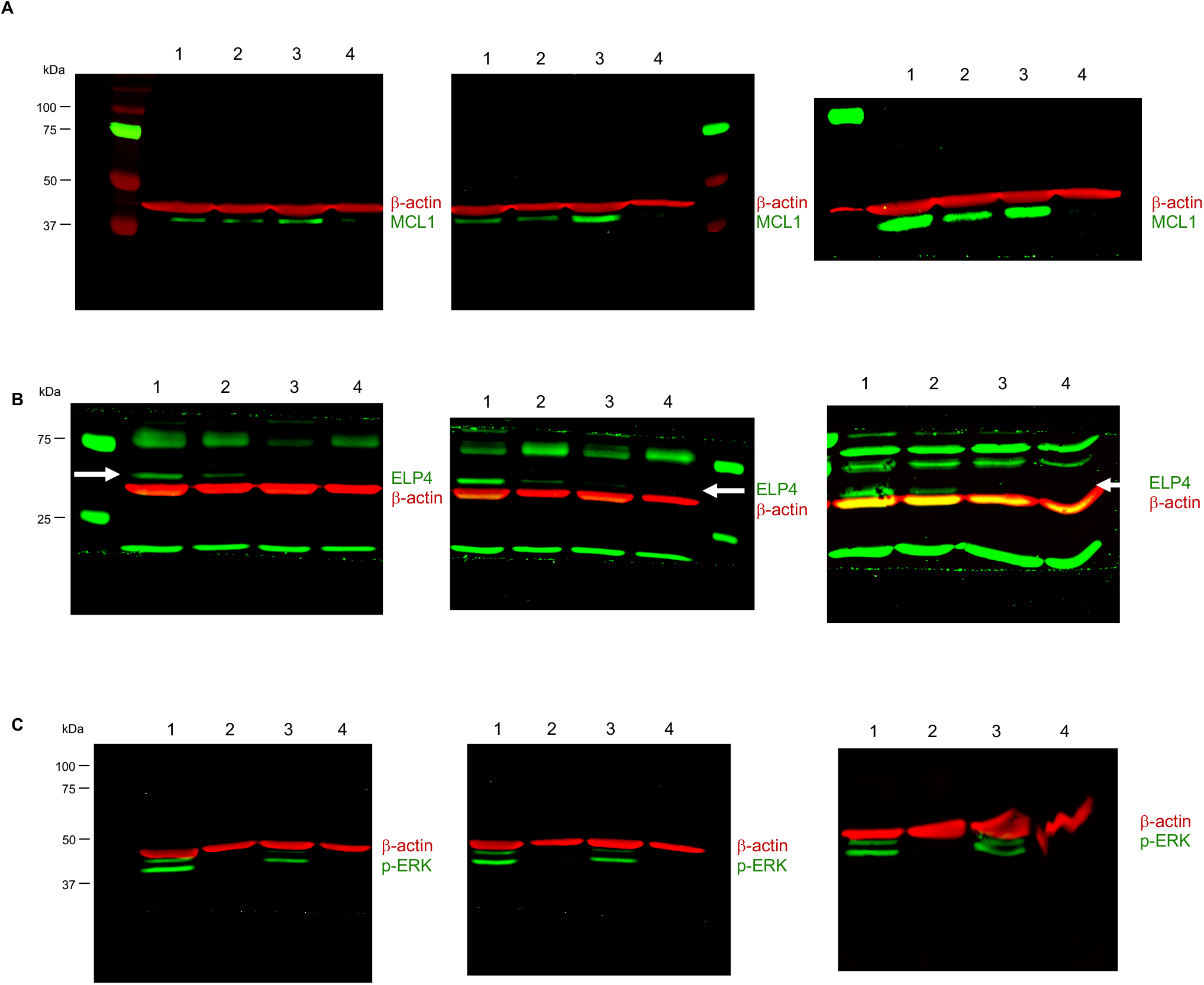
ELP complex facilitates expression of MCL1 following erlotinib exposure. **(A-C)** Western blots for MCL1 (A), ELP4 (B), and phosphorylated ERK (p-ERK, C). Beta-actin shown as a loading control (β-actin). Three independent biological replicates shown. Data are 1: scrambled RNA, untreated; 2: scrambled RNA + 10 μM erlotinib; 3: ELP4 KD, untreated; 4: ELP4 KD + 10 μM erlotinib. In panel (B), white arrow marks the expected size/band for ELP4. See also Fig. 3.

**Fig. S6:**
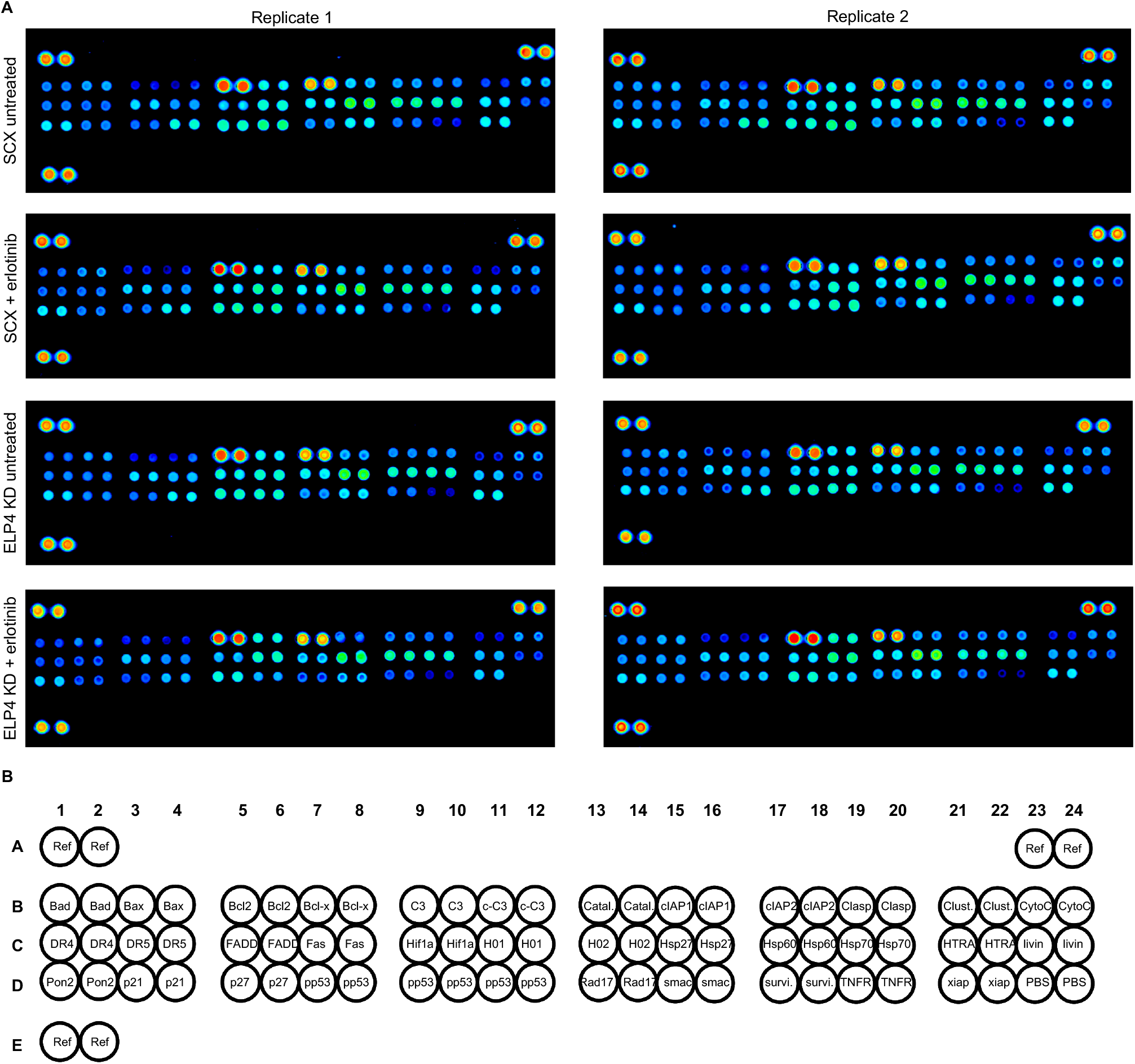
Protein level changes within key apoptotic regulatory proteins following erlotinib exposure. **(A-B)** Proteome profiler human apoptosis array. Data shown for BT20 cells with scrambled RNA (SCX) or ELP4 targeted siRNA (ELP4 KD), each with or without 10 μM erlotinib. Two independent biological replicates of each shown. **(B)** Apoptosis array analysis key.

**Fig. S7:**
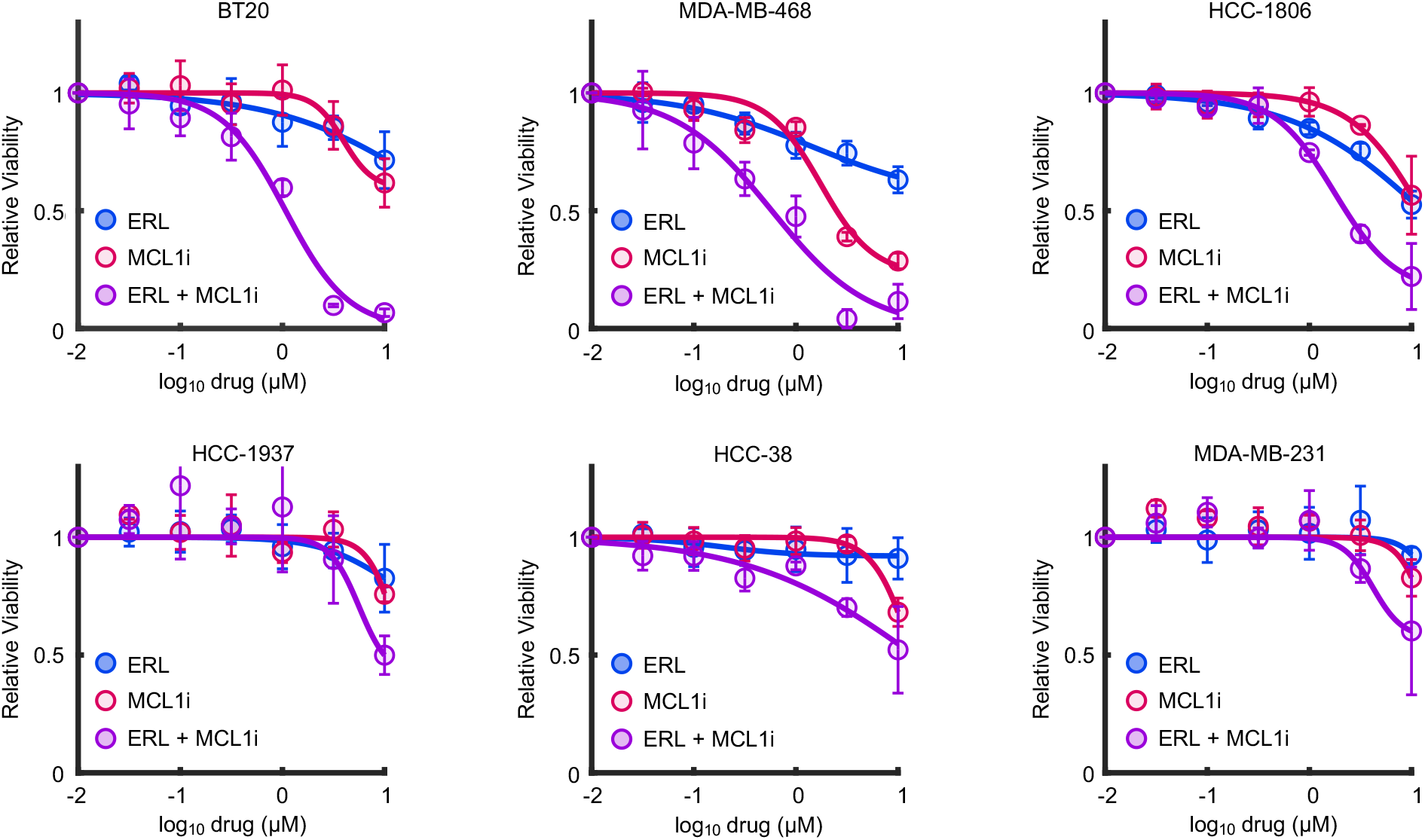
MCL1 inhibition synergistically enhances sensitivity to erlotinib in a panel TNBC cells. Sensitivity of TNBC cells to erlotinib (ERL), S63845 (MCL1i), or a combination of these two drugs (ERL + MCL1i). Drugs tested across 7 half-log dilutions. For drug combinations, ERL and MCL1i were added at a fixed 1:1 equimolar drug ratio. Relative Viability measured 72 hours after drug exposure using a SYTOX based death assay. Data are mean +/− s.d. of 3 independent biological replicates. See also Fig. 4E and F.

## REFERENCES

1. S. V. Sharma, J. Settleman, Oncogene addiction: setting the stage for molecularly targeted cancer therapy, Gene Dev 21, 3214–3231 (2007).

2. I. B. Weinstein, M. Begemann, P. Zhou, E. K. Han, A. Sgambato, Y. Doki, N. Arber, M. Ciaparrone, H. Yamamoto, Disorders in cell circuitry associated with multistage carcinogenesis: exploitable targets for cancer prevention and therapy., Clin Cancer Res Official J Am Assoc Cancer Res 3, 2696–702 (1997).

3. R. Pagliarini, W. Shao, W. R. Sellers, Oncogene addiction: pathways of therapeutic response, resistance, and road maps toward a cure, Embo Rep 16, 280–296 (2015).

4. R. B. Corcoran, H. Ebi, A. B. Turke, E. M. Coffee, M. Nishino, A. P. Cogdill, R. D. Brown, P. D. Pelle, D. Dias-Santagata, K. E. Hung, K. T. Flaherty, A. Piris, J. A. Wargo, J. Settleman, M. Mino-Kenudson, J. A. Engelman, EGFR-Mediated Reactivation of MAPK Signaling Contributes to Insensitivity of BRAF-Mutant Colorectal Cancers to RAF Inhibition with Vemurafenib, Cancer Discov 2, 227–235 (2012).

5. H. A. Wahba, H. A. El-Hadaad, Current approaches in treatment of triple-negative breast cancer, Cancer Biology Medicine 12, 106–116 (2015).

6. P. Savage, A. Blanchet-Cohen, T. Revil, D. Badescu, S. M. I. Saleh, Y.-C. Wang, D. Zuo, L. Liu, N. R. Bertos, V. Munoz-Ramos, M. Basik, K. Petrecca, J. Asselah, S. Meterissian, M.-C. Guiot, A. Omeroglu, C. L. Kleinman, M. Park, J. Ragoussis, A Targetable EGFR-Dependent Tumor-Initiating Program in Breast Cancer, Cell Reports 21, 1140–1149 (2017).

7. B. Corkery, J. Crown, M. Clynes, N. O’donovan, Epidermal growth factor receptor as a potential therapeutic target in triple-negative breast cancer, Ann Oncol 20, 6 (2009).

8. T. Sun, N. Aceto, K. L. Meerbrey, J. D. Kessler, C. Zhou, I. Migliaccio, D. X. Nguyen, N. N. Pavlova, M. Botero, J. Huang, R. J. Bernardi, E. Schmitt, G. Hu, M. Z. Li, N. Dephoure, S. P. Gygi, M. Rao, C. J. Creighton, S. G. Hilsenbeck, C. A. Shaw, D. Muzny, R. A. Gibbs, D. A. Wheeler, C. K. Osborne, R. Schiff, M. Bentires-Alj, S. J. Elledge, T. F. Westbrook, Activation of multiple proto-oncogenic tyrosine kinases in breast cancer via loss of the PTPN12 phosphatase., Cell 144, 703–718 (2011).

9. T. O. Nielsen, F. D. Hsu, K. Jensen, M. Cheang, G. Karaca, Z. Hu, T. Hernandez-Boussard, C. Livasy, D. Cowan, L. Dressler, L. A. Akslen, J. Ragaz, A. M. Gown, C. B. Gilks, M. van de Rijn, C. M. Perou, Immunohistochemical and Clinical Characterization of the Basal-Like Subtype of Invasive Breast Carcinoma, Clin Cancer Res 10, 5367–5374 (2004).

10. V. Secq, J. Villeret, F. Fina, M. Carmassi, X. Carcopino, S. Garcia, I. Metellus, L. Boubli, J. Iovanna, C. Charpin, Triple negative breast carcinoma EGFR amplification is not associated with EGFR, Kras or ALK mutations, Brit J Cancer 110, 1045–1052 (2014).

11. R. Sordella, D. W. Bell, D. A. Haber, J. Settleman, Gefitinib-sensitizing EGFR mutations in lung cancer activate anti-apoptotic pathways., Science 305, 1163–1167 (2004).

12. L. V. Sequist, B. A. Waltman, D. Dias-Santagata, S. Digumarthy, A. B. Turke, P. Fidias, K. Bergethon, A. T. Shaw, S. Gettinger, A. K. Cosper, S. Akhavanfard, R. S. Heist, J. Temel, J. G. Christensen, J. C. Wain, T. J. Lynch, K. Vernovsky, E. J. Mark, M. Lanuti, A. J. Iafrate, M. Mino-Kenudson, J. A. Engelman, Genotypic and histological evolution of lung cancers acquiring resistance to EGFR inhibitors., Sci Transl Med 3, 75ra26–75ra26 (2011).

13. L. A. Carey, H. S. Rugo, P. K. Marcom, E. L. Mayer, F. J. Esteva, C. X. Ma, M. C. Liu, A. M. Storniolo, M. F. Rimawi, A. Forero-Torres, A. C. Wolff, T. J. Hobday, A. Ivanova, W. K. Chiu, M. Ferraro, E. Burrows, P. S. Bernard, K. A. Hoadley, C. M. Perou, E. P. Winer, TBCRC 001: Randomized Phase II Study of Cetuximab in Combination With Carboplatin in Stage IV Triple-Negative Breast Cancer, J Clin Oncol 30, 2615–2623 (2012).

14. R. Richards, H. R. Schwartz, M. S. Stewart, P. Cruz-Gordillo, M. E. Honeywell, A. J. Joyce, B. D. Landry, M. J. Lee, Drug Combination Antagonism and Single Agent Dominance Result from Differences in Death Activation Kinetics, Biorxiv , 805093 (2019).

15. M. Hafner, M. Niepel, M. Chung, P. K. Sorger, Growth rate inhibition metrics correct for confounders in measuring sensitivity to cancer drugs, Nat Methods 13, 1–11 (2016).

16. M. Ono, A. Hirata, T. Kometani, M. Miyagawa, S. Ueda, H. Kinoshita, T. Fujii, M. Kuwano, Sensitivity to gefitinib (Iressa, ZD1839) in non-small cell lung cancer cell lines correlates with dependence on the epidermal growth factor (EGF) receptor/extracellular signal-regulated kinase 1/2 and EGF receptor/Akt pathway for proliferation., Mol Cancer Ther 3, 465–72 (2004).

17. G. C. Forcina, M. Conlon, A. Wells, J. Y. Cao, S. J. Dixon, Systematic Quantification of Population Cell Death Kinetics in Mammalian Cells, Cell Syst 4, 1–18 (2017).

18. H. R. Schwartz, R. Richards, R. E. Fontana, A. J. Joyce, M. E. Honeywell, M. J. Lee, Drug GRADE: an integrated analysis of population growth and cell death reveals drug-specific and cancer subtype-specific response profiles, bioRxiv (2020), doi:10.1101/2020.02.26.966689.

19. D. T. Worster, T. Schmelzle, N. L. Solimini, E. S. Lightcap, B. Millard, G. B. Mills, J. S. Brugge, J. G. Albeck, Akt and ERK control the proliferative response of mammary epithelial cells to the growth factors IGF-1 and EGF through the cell cycle inhibitor p57Kip2., Sci Signal 5, ra19–ra19 (2012).

20. J. Luo, N. L. Solimini, S. J. Elledge, Principles of cancer therapy: oncogene and non-oncogene addiction., Cell 136, 823–837 (2009).

21. G. D. Guler, C. A. Tindell, R. Pitti, C. Wilson, K. Nichols, T. K. Cheung, H.-J. Kim, M. Wongchenko, Y. Yan, B. Haley, T. Cuellar, J. Webster, N. Alag, G. Hegde, E. Jackson, T. L. Nance, P. G. Giresi, K.-B. Chen, J. Liu, S. Jhunjhunwala, J. Settleman, J.-P. Stephan, D. Arnott, M. Classon, Repression of Stress-Induced LINE-1 Expression Protects Cancer Cell Subpopulations from Lethal Drug Exposure, Cancer Cell 32, 221–237.e13 (2017).

22. A. Subramanian, P. Tamayo, V. K. Mootha, S. Mukherjee, B. L. Ebert, M. A. Gillette, A. Paulovich, S. L. Pomeroy, T. R. Golub, E. S. Lander, J. P. Mesirov, Gene set enrichment analysis: a knowledge-based approach for interpreting genome-wide expression profiles., Proc National Acad Sci 102, 15545–15550 (2005).

23. S. Kobayashi, T. Shimamura, S. Monti, U. Steidl, C. J. Hetherington, A. M. Lowell, T. Golub, M. Meyerson, D. G. Tenen, G. I. Shapiro, B. Halmos, Transcriptional Profiling Identifies Cyclin D1 as a Critical Downstream Effector of Mutant Epidermal Growth Factor Receptor Signaling, Cancer Res 66, 11389–11398 (2006).

24. O. Shalem, N. E. Sanjana, E. Hartenian, X. Shi, D. A. Scott, T. S. Mikkelsen, D. Heckl, B. L. Ebert, D. E. Root, J. G. Doench, F. Zhang, Genome-scale CRISPR-Cas9 knockout screening in human cells., Science 343, 84–87 (2014).

25. K. Imkeller, G. Ambrosi, M. Boutros, W. Huber, gscreend: modelling asymmetric count ratios in CRISPR screens to decrease experiment size and improve phenotype detection, Genome Biol 21, 53 (2020).

26. O. Parnas, M. Jovanovic, T. M. Eisenhaure, R. H. Herbst, A. Dixit, C. J. Ye, D. Przybylski, R. J. Platt, I. Tirosh, N. E. Sanjana, O. Shalem, R. Satija, R. Raychowdhury, P. Mertins, S. A. Carr, F. Zhang, N. Hacohen, A. Regev, A Genome-wide CRISPR Screen in Primary Immune Cells to Dissect Regulatory Networks, Cell 162, 675–686 (2015).

27. W. Li, H. Xu, T. Xiao, L. Cong, M. I. Love, F. Zhang, R. A. Irizarry, J. S. Liu, M. Brown, X. S. Liu, MAGeCK enables robust identification of essential genes from genome-scale CRISPR/Cas9 knockout screens, Genome Biol 15, 819–12 (2014).

28. M. Colic, G. Wang, M. Zimmermann, K. Mascall, M. McLaughlin, L. Bertolet, W. F. Lenoir, J. Moffat, S. Angers, D. Durocher, T. Hart, Identifying chemogenetic interactions from CRISPR screens with drugZ, Genome Med 11, 52 (2019).

29. T. Hart, J. Moffat, BAGEL: a computational framework for identifying essential genes from pooled library screens, Bmc Bioinformatics 17, 164 (2016).

30. B. Luo, H. W. Cheung, A. Subramanian, T. Sharifnia, M. Okamoto, X. Yang, G. Hinkle, J. S. Boehm, R. Beroukhim, B. A. Weir, C. Mermel, D. A. Barbie, T. Awad, X. Zhou, T. Nguyen, B. Piqani, C. Li, T. R. Golub, M. Meyerson, N. Hacohen, W. C. Hahn, E. S. Lander, D. M. Sabatini, D. E. Root, Highly parallel identification of essential genes in cancer cells., Proc National Acad Sci 105, 20380–20385 (2008).

31. T. Hart, M. Chandrashekhar, M. Aregger, Z. Steinhart, K. R. Brown, G. MacLeod, M. Mis, M. Zimmermann, A. Fradet-Turcotte, S. Sun, P. Mero, P. Dirks, S. Sidhu, F. P. Roth, O. S. Rissland, D. Durocher, S. Angers, J. Moffat, High-Resolution CRISPR Screens Reveal Fitness Genes and Genotype-Specific Cancer Liabilities, Cell 163, 1515–1526 (2015).

32. G. Wang, M. Zimmermann, K. Mascall, W. F. Lenoir, J. Moffat, S. Angers, D. Durocher, T. Hart, Identifying drug-gene interactions from CRISPR knockout screens with drugZ, Biorxiv , 1–16 (2017).

33. G. Otero, J. Fellows, Y. Li, T. de Bizemont, A. M. G. Dirac, C. M. Gustafsson, H. Erdjument-Bromage, P. Tempst, J. Q. Svejstrup, Elongator, a Multisubunit Component of a Novel RNA Polymerase II Holoenzyme for Transcriptional Elongation, Mol Cell 3, 109–118 (1999).

34. J.-H. Kim, W. S. Lane, D. Reinberg, Human Elongator facilitates RNA polymerase II transcription through chromatin, Proc National Acad Sci 99, 1241–1246 (2002).

35. B. Huang, M. J. O. Johansson, A. S. BystrÖm, An early step in wobble uridine tRNA modification requires the Elongator complex, Rna 11, 424–436 (2005).

36. J. Lorent, E. P. Kusnadi, V. Hoef, R. J. Rebello, M. Leibovitch, J. Ristau, S. Chen, M. G. Lawrence, K. J. Szkop, B. Samreen, P. Balanathan, F. Rapino, P. Close, P. Bukczynska, K. Scharmann, I. Takizawa, G. P. Risbridger, L. A. Selth, S. A. Leidel, Q. Lin, I. Topisirovic, O. Larsson, L. Furic, Translational offsetting as a mode of estrogen receptor α-dependent regulation of gene expression, Embo J 38(2019), doi:10.15252/embj.2018101323.

37. F. Rapino, S. Delaunay, F. Rambow, Z. Zhou, L. Tharun, P. D. Tullio, O. Sin, K. Shostak, S. Schmitz, J. Piepers, B. Ghesquière, L. Karim, B. Charloteaux, D. Jamart, A. Florin, C. Lambert, A. Rorive, G. Jerusalem, E. Leucci, M. Dewaele, M. Vooijs, S. A. Leidel, M. Georges, M. Voz, B. Peers, R. Büttner, J.-C. Marine, A. Chariot, P. Close, Codon-specific translation reprogramming promotes resistance to targeted therapy, Nature 558, 605–609 (2018).

38. M. Dewez, F. Bauer, M. Dieu, M. Raes, J. Vandenhaute, D. Hermand, The conserved Wobble uridine tRNA thiolase Ctu1-Ctu2 is required to maintain genome integrity., P Natl Acad Sci Usa 105, 5459–64 (2008).

39. T. Karlsborn, H. Tükenmez, A. K. M. F. Mahmud, F. Xu, H. Xu, A. S. Byström, Elongator, a conserved complex required for wobble uridine modifications in Eukaryotes, Rna Biol 11, 1519–1528 (2015).

40. D. D. Nedialkova, S. A. Leidel, Optimization of Codon Translation Rates via tRNA Modifications Maintains Proteome Integrity, Cell 161, 1606–1618 (2015).

41. K. M. Kozopas, T. Yang, H. L. Buchan, P. Zhou, R. W. Craig, MCL1, a gene expressed in programmed myeloid cell differentiation, has sequence similarity to BCL2., Proc National Acad Sci 90, 3516–3520 (1993).

42. A. Kotschy, Z. Szlavik, J. Murray, J. Davidson, A. L. Maragno, G. L. Toumelin-Braizat, M. Chanrion, G. L. Kelly, J.-N. Gong, D. M. Moujalled, A. Bruno, M. Csekei, A. Paczal, Z. B. Szabo, S. Sipos, G. Radics, A. Proszenyak, B. Balint, L. Ondi, G. Blasko, A. Robertson, A. Surgenor, P. Dokurno, I. Chen, N. Matassova, J. Smith, C. Pedder, C. Graham, A. Studeny, G. Lysiak-Auvity, A.-M. Girard, F. Gravé, D. Segal, C. D. Riffkin, G. Pomilio, L. C. A. Galbraith, B. J. Aubrey, M. S. Brennan, M. J. Herold, C. Chang, G. Guasconi, N. Cauquil, F. Melchiore, N. Guigal-Stephan, B. Lockhart, F. Colland, J. A. Hickman, A. W. Roberts, D. C. S. Huang, A. H. Wei, A. Strasser, G. Lessene, O. Geneste, The MCL1 inhibitor S63845 is tolerable and effective in diverse cancer models, Nature 538, 477–482 (2016).

43. C. T. Meyer, D. J. Wooten, B. B. Paudel, J. Bauer, K. N. Hardeman, D. Westover, C. M. Lovly, L. A. Harris, D. R. Tyson, V. Quaranta, Quantifying Drug Combination Synergy along Potency and Efficacy Axes, Cell Syst 8, 1–44 (2019).

44. D. Russ, R. Kishony, Additivity of inhibitory effects in multidrug combinations, Nat Microbiol 3, 1–9 (2018).

45. C. T. Meyer, D. J. Wooten, C. F. Lopez, V. Quaranta, Charting the Fragmented Landscape of Drug Synergy., Trends Pharmacol Sci (2020), doi:10.1016/j.tips.2020.01.011.

46. T. Yang, K. M. Kozopas, R. W. Craig, The intracellular distribution and pattern of expression of Mcl-1 overlap with, but are not identical to, those of Bcl-2., J Cell Biology 128, 1173–1184 (1995).

47. A. M. Domina, J. A. Vrana, M. A. Gregory, S. R. Hann, R. W. Craig, MCL1 is phosphorylated in the PEST region and stabilized upon ERK activation in viable cells, and at additional sites with cytotoxic okadaic acid or taxol, Oncogene 23, 5301–5315 (2004).

48. J.-M. Wang, J.-R. Chao, W. Chen, M.-L. Kuo, J. J.-Y. Yen, H.-F. Yang-Yen, The Antiapoptotic Gene mcl-1 Is Up-Regulated by the Phosphatidylinositol 3-Kinase/Akt Signaling Pathway through a Transcription Factor Complex Containing CREB, Mol Cell Biol 19, 6195–6206 (1999).

